# Unbiased integration of single cell transcriptome replicates

**DOI:** 10.1101/2021.05.05.442380

**Authors:** Martin Loza Lopez, Shunsuke Teraguchi, Daron M. Standley, Diego Diez

**Affiliations:** Immunology Frontier Research Center, Osaka University, Suita, 565-0871, Japan; Faculty of Data Science, Shiga University, Hikone, 522-8522, Japan; Research Institute for Microbial Diseases, Osaka University, Suita, 565-0871, Japan

## Abstract

Single cell transcriptomic approaches are becoming mainstream, with replicate experiments commonly performed with the same single cell technology. Methods that enable integration of these datasets by removing batch effects while preserving biological information are required for unbiased data interpretation. Here we introduce Canek for this purpose. Canek leverages information from mutual nearest neighbor to combine local linear corrections with cell-specific non-linear corrections within a fuzzy logic framework. Using a combination of real and synthetic datasets, we show that Canek corrects batch effects while introducing the least amount of bias compared with competing methods. Canek is computationally efficient and can easily integrate thousands of single-cell transcriptomes from replicated experiments.

## Introduction

Single cell sequencing technologies allow quantification of RNA expression levels within a given cell with unprecedented resolution [1]. However, these approaches require integration over multiple observations to increase signal to noise. In such efforts, the true biological signal can become distorted. Even the most skilled operator using the same instrument will tend to observe systematic differences in replicates. Although such batch effects are well-known, they do not result from a single cause and thus are difficult to define or correct [2].

Many methods to integrate single cell datasets obtained from the same tissues using different technologies have been introduced [3]. One of the pioneering techniques is the so-called Mutual Nearest Neighbors (MNN) correction method [4]. In this method, MNN pairs are used to identify corresponding cells in two different batches. A pair-specific correction vector is then defined as the difference between the expression profiles of the cells from each MNN pair. The correction vectors are then weighted to smooth the corrections between adjacent cells. Subsequently, other tools have been developed that use MNNs to integrate batches [5–7]. One popular method, implemented in the Seurat R package, finds MNN pairs in a correlated space using canonical correlation analysis (CCA) [5]. The identified pairs are used as “anchors” to correct batch effects. Another interesting approach, Harmony, iteratively removes batch effects by clustering in a low dimensional space [8]. LIGER applies a similar clustering approach by segmenting cells using a shared factor neighborhood graph under a low dimensional space defined with an integrative non-negative matrix factorization method [9].

In a comprehensive benchmark of 14 batch correction methods, including the ones above, the methods were tested under different scenarios to quantify the effects of data acquired by different technologies, use of dissimilar cells, data size, numbers of batches and simulated biases [3]. The three top-scoring methods were Harmony, Liger, and Seurat. However, the authors found that each method performed differently on each test, with no obviously superior method [3]. Another benchmark done on atlas-level datasets found that the best integration approach strongly depended on the target task [10]. This ambiguity makes best practices for integration of batches from replicated experiments unclear. An important question is how much bias such methods introduce and which of the methods is best suited for this task. To address these issues, we introduce Canek. Canek operates on two levels: it assumes mostly linear batch effects within a cluster of similar cells but allows non-linear corrections between different clusters of cells in a pair of datasets. This allows Canek to efficiently integrate single cell transcriptomes from replicated experiments while introducing minimal bias, thus preserving biologically relevant information.

## Results

### Overview of Canek

Canek corrects multiple batches by integrating pairs of batches sequentially. The dataset pairs that are input to Canek are denoted reference batch and query batch (Figure 1a). Then the integrated dataset becomes the reference to integrate the following batch. *Batch effect observations are defined* using *mutual nearest neighbors* (MNN) [4] and groups of similar cells are identified from the *query batch* using *clustering* (Figure 1b). Canek estimates a *correction vector* for each cluster using the median gene expression differences between cells in each cluster of the query batch and the corresponding cells in the reference as identified by MNN (arrows on Figure 1c). The correction vector can thus be used to remove the batch effect from each cluster in the query batch. In this linear correction, the same correction is applied to all the cells in the cluster (Figure 1c). Subsequently, Canek performs a *non-linear correction* by calculating a cell-specific transformation using fuzzy logic. This is done by defining a minimum spanning tree among clusters and then smoothing the transitions between the correction vectors (Figure 1d). Using a combination of real and simulated data, we show that, Canek exhibits unbiased corrections of single cell transcriptome data.

**Figure 1.**
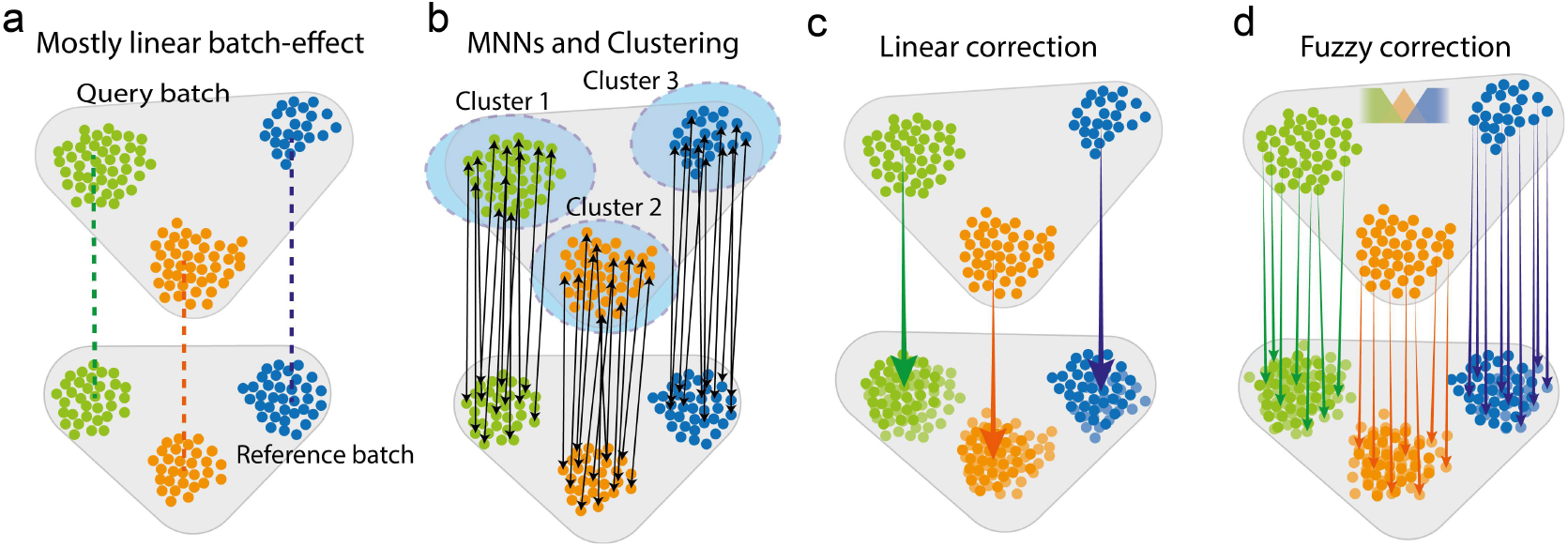
Overview of Canek workflow. **a)** *Canek starts with a* reference batch and query batch, assuming a predominantly linear batch effect. **b)** Cell clusters are defined on the query batch and MNN pairs (arrows) are used to define batch effect observations. **c)** The MNN pairs from each cluster are used to estimate cluster specific correction vectors. These vectors can be used to correct the batch effect or, **d)** a non-linear correction can be applied by calculating cell-specific correction vectors using fuzzy logic.

### Canek successfully corrects batch effects in a Jurkat/293T mixture dataset

An example where batch effects are clearly visible is the mixture of cells used to demonstrate the 10x chromium sequencing technology [11]. The dataset consists of three batches: one containing only 293T HEK (Human Embryonic Kidney) cells, another containing only Jurkat cells (immortalized human T lymphocytes), and a third comprised of a 50:50 mixture of 293T and Jurkat cells [11]. Principal component analysis (PCA) of the Uncorrected dataset is shown in Figure 2a. Looking at cell-specific markers we can see there is a cluster of cells expressing XIST (293T cells) and two clusters of cells expressing CD3D (Jurkat cells). While the cluster of 293T cells shows mixing of cells from both batches, the two clusters of Jurkat cells show batch specific distributions, suggesting an unknown systematic bias. We used different integration methods and assessed their ability to correct the systematic differences in the Jurkat cell data without introducing additional bias. To this end, we applied batch correction using Canek and 8 state-of-the-art methods: Combat, ComBat-seq, Harmony, Liger, MNN, Scanorama, scMerge, and Seurat [4, 5, 7–9, 12–14]. Both Canek and MNN corrected the batch effect and enabled the identification of the expected cell population clusters (Figure 2b,c). However, other methods, including Combat and Seurat, resulted in incorrect mixing of cell populations (Figure 2d, e). The results for all methods are shown in Supplementary Figure 1 for PCA, and Supplementary Figure 2 for Uniform Manifold Approximation and Projection (UMAP) [15] plots.

**Figure 2.**
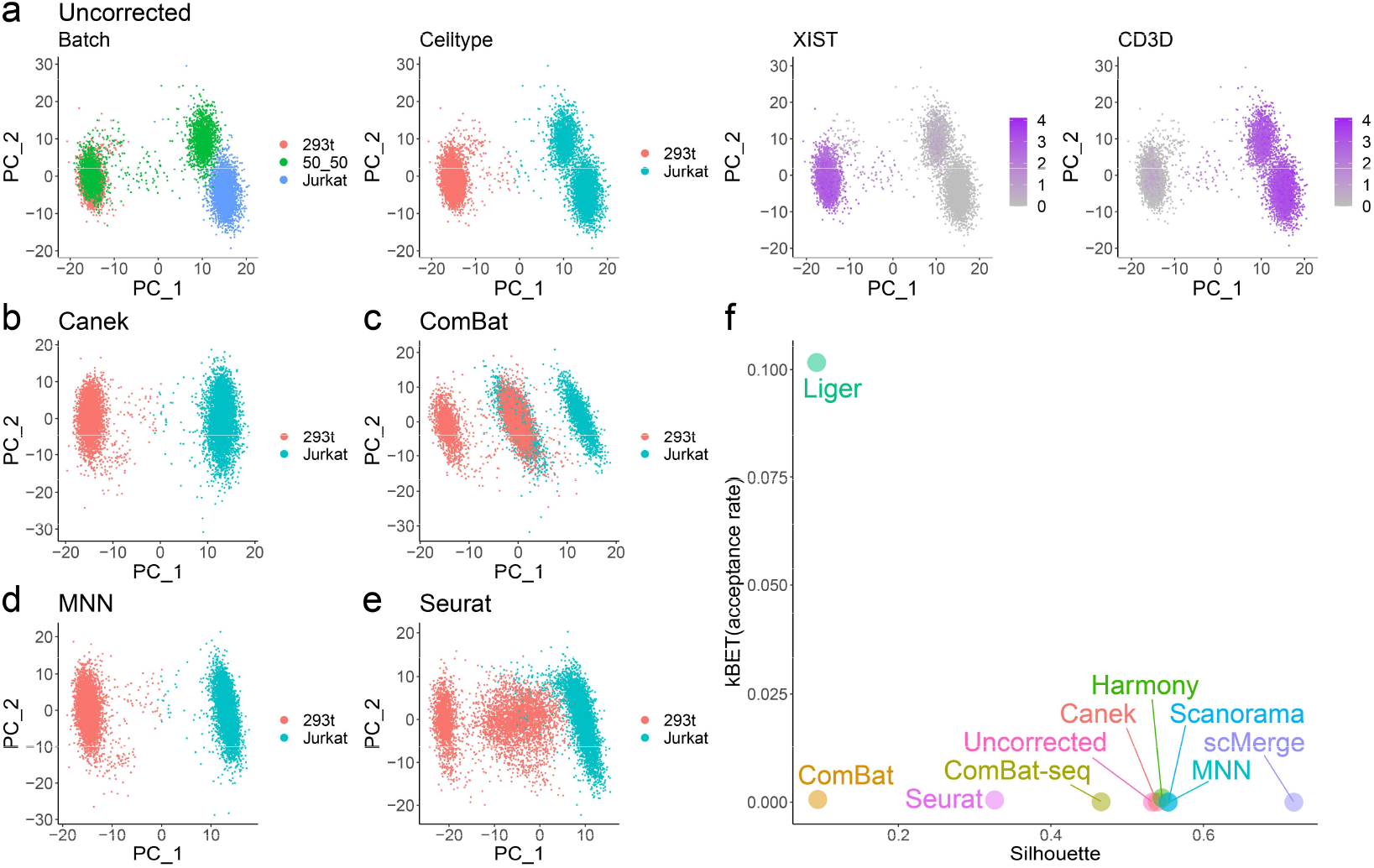
Batch effect correction methods may incorrectly mix dissimilar cell types. Batch effect correction of three batches, two containing pure Jurkat and HEK293T cells, and one with a 50:50 mix of Jurkat and HEK293T cells. **a)** Jurkat and HEK293T cells are characterized by the expression of CD3D and XIST genes respectively. Before correction Jurkat cells grouped by batch. **b-e)** *show the results of ba*tch effect correction using Canek, Combat, MNN and Seurat. Canek and MNN correctly integrated the Jurkat cells. Combat and Seurat incorrectly mixed Jurkat and HEK293T cells.

To evaluate the performance of batch correction we computed kBET and Silhouette scores for the Uncorrected and corrected datasets (Figure 2f). We chose the kBET metric to estimate the mixing of batches after correction, while the Silhouette metric enabled us to assess the preservation of the original cell clusters. Most methods show similar values of kBET, indicating similar levels of mixing. Methods that successfully integrated the batches while preserving cell populations had higher values in the Silhouette score. Canek, Harmony, MNN, and Scanorama all have similar values, and lead to visually successful integrations as seen in the PCA and UMAP plots. Combat, ComBat-seq and Seurat resulted in good integration but different levels of success in preserving the cell populations. Liger showed very different behavior (high kBET and low Silhouette), suggesting excessive mixing while not preserving cell populations. This agrees with the PCA/UMAP plots in Supplementary Figures 1 and 2. These results show that Canek was able to successfully identify and correct the local batch effect while preserving biologically meaningful cell type differences.

### Evaluation of correction bias

As shown above, batch correction methods can introduce biases in the data that disturb the biological information or alter the structure of cell populations [10, 16]. As single-cell genomics technologies become mainstream, more laboratories will perform experiments under different conditions with biological replicates obtained using a common technology. In this scenario, integration of datasets with minimal impact on cell phenotype is essential.

We define batch correction bias as undesired correction that may alter the original biological signal. To quantify how much bias correction methods introduce, we use a pseudo-batch approach (shown schematically in Figure 3a). Starting from a single dataset we identified clusters to define cell populations. Then we generated two pseudo-batches by sampling cells without replacement. Each pseudo-batch preserves the information about the original cell populations (clusters). Because no modifications to the original expression values were introduced during the sampling process, the batch effect between the two pseudo-batches is effectively zero. We assume that batch correction methods should not correct in this scenario since no batch effect exists, and identification of clusters from the integrated batches should preserve the clusters obtained from the original (Uncorrected) dataset.

**Figure 3.**
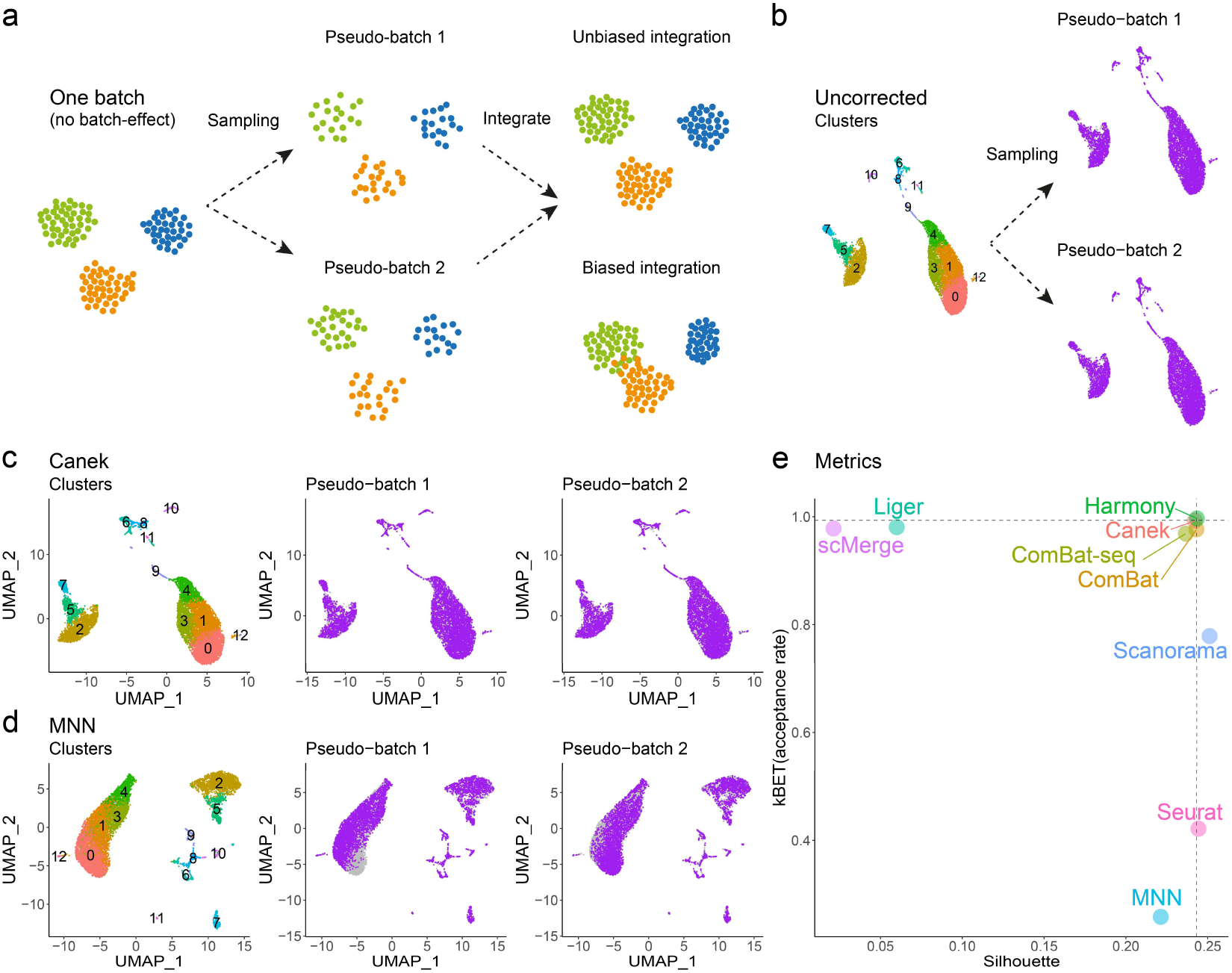
Correction methods may introduce biases. a) Strategy for pseudo-batch generation: Starting from a single dataset with identified cell populations (clusters), we sample without replacement to generate two pseudo-batches. Then, we integrate these pseudo-batches and check whether the integration introduced biases comparing the result with the original dataset. b) Pseudo-batch generation using the spleen dataset from Tabula Muris. c) Canek integrated the pseudo-batches without introducing biases to the cell distribution. e) MNN integration led to uneven distribution of cells from the pseudo-batches in the UMAP plot. e) Using kBET and Silhouette metrics, the mixing among batches and the cluster preservation were evaluated. The optimal scores from the Uncorrected data are shown as dashed lines, while the scores from the correction methods are indicated as colored points. Unbiased methods are those whose metrics are closest to the intersection of the gray lines.

We applied this strategy to the droplet spleen dataset from Tabula Muris [17]. In Figure 3b the first UMAP plot shows the original dataset with cell clusters indicated with colors. The next two UMAP plots show the cells from the two pseudo-batches obtained after sampling. We applied batch correction to these two pseudo-batches. To quantify the introduced bias, we computed kBET and Silhouette scores for the Uncorrected and corrected datasets. Since there was no batch effect, the scores for the Uncorrected dataset corresponded to the optimal values.

Figure 3c shows that Canek integration resulted in even distribution of the cells from both batches and cluster distribution that resembled the original dataset. Figure 3d shows that MNN failed to completely integrate this dataset, with uneven distribution of cells from the pseudo-batches. In Supplementary Figure 3 we show UMAP plots with results for each of the evaluated methods. Although in some cases it was trivial to identify differences with the Uncorrected dataset due to obvious changes in cell distributions, it was not always easy to evaluate the relative performance. To do so, we calculated kBET and Silhouette scores and compared them with those obtained from the Uncorrected dataset, which represented the optimal values. Figure 3e shows the scores for kBET and Silhouette metrics obtained from this experiment. In this plot, the dashed lines indicate the scores for the Uncorrected dataset, with the crossing point representing the optimal value. Canek, Combat, ComBat-seq and Harmony resulted in scores very close to the optimal value. To estimate the variability of the results due to pseudo-batch sampling, we repeated this experiment 10 times. Supplementary Figure 4 shows that Canek obtained scores closest to the values of the Uncorrected dataset, demonstrating that it introduced the least amount of bias when no batch effect was present.

### Evaluation of integration in simulated data

To estimate the ability to correct batch effects when the effect is known exactly, we compared Canek with other methods using simulated data. We simulated three batches with shared cell types using the splatter package [18] from which we can obtain an integrated dataset to use as a gold standard (GS). Batch 1 was composed of two shared and one unique cell type, whereas batches 2 and 3 had one shared and one unique cell type (see Table 2 for a complete description). Figures 4a and 4b show UMAP plots from the GS and the Uncorrected dataset, respectively. Figure 4c shows kBET and Silhouette scores from the GS (cross of dashed lines), Uncorrected data, and integrated datasets. We expected the best correction methods to be close to the metrics from GS. These results show that Canek scores were closest to those of the GS. This is consistent with the UMAP plot shown in Figure 4d, where Canek corrected the differences among shared cell types. Interestingly, Harmony returned scores very close to the Uncorrected data, suggesting that it performed almost no correction, consistent with the UMAP plot in Figure 4e.

**Figure 4.**
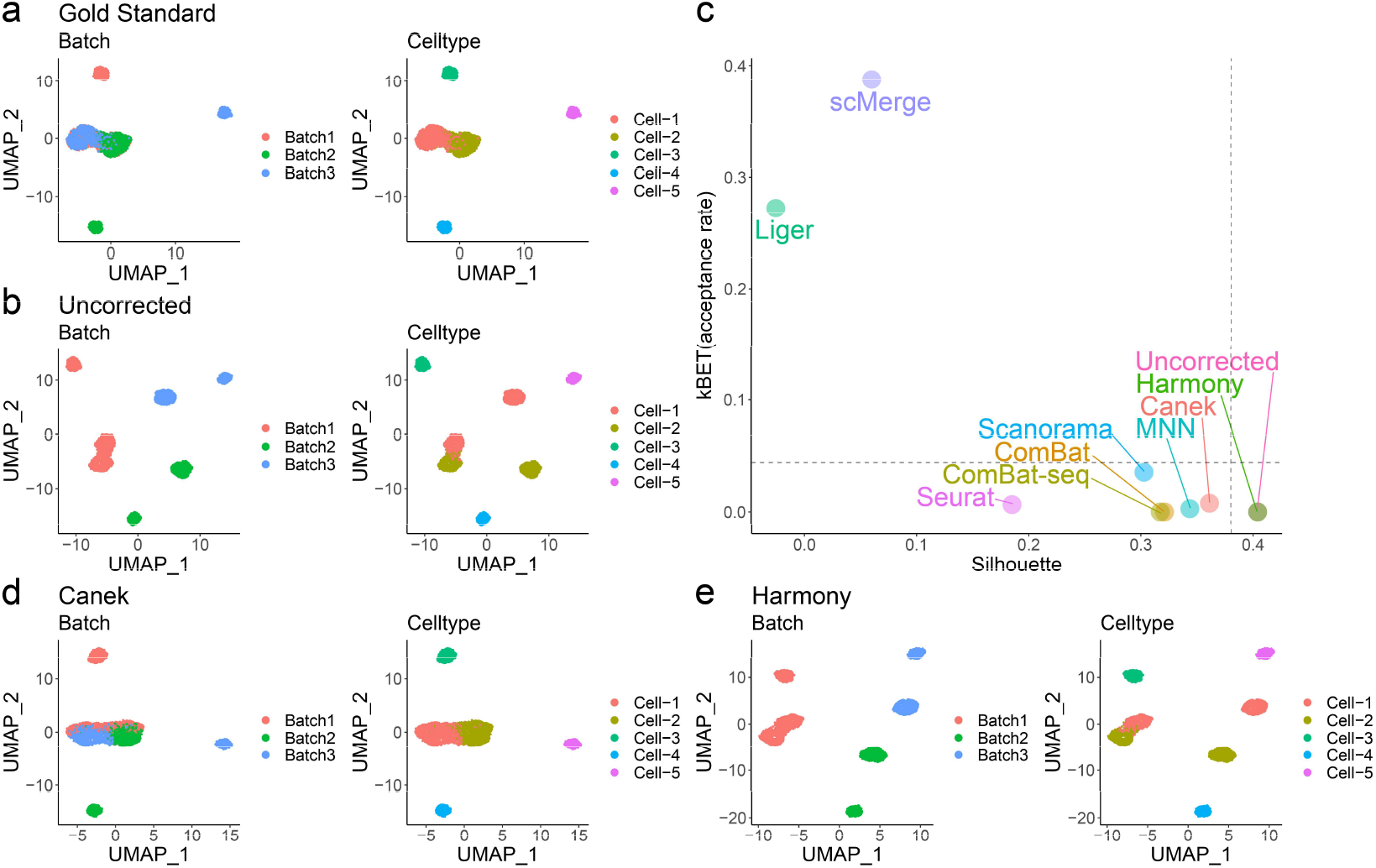
Batch effect correction on simulated data with a known gold standard. Three batches were simulated to test the integration methods in a scenario with a known gold standard **a)**. The gold standard shows the UMAP plot for the batches without batch effect. Two cell types (Cell-1 and Cell-2) are shared among different batches. **b)** The Uncorrected dataset shows batch-specific differences in cells of the same type. **c)** kBET and Silhouette metrics for Uncorrected, Gold Standard (dashed lines) and the 9 evaluated methods. Canek shows scores closest to the Gold Standard. **d)** UMAP plot shows that Canek correctly integrated the shared cell types while maintaining the identity of non-shared ones. **e)** Harmony correction failed to integrate cells from the same type.

**Table 1.** Batch effect correction methods used.

| Method | Batch effect corrected output | Package version |
| --- | --- | --- |
| Seurat | Normalized gene expression matrix | Seurat version 3.2.2 [5] |
| MNN | Normalized gene expression matrix | Bioconductor’s batchelor version 1.6.2 [4] |
| Scanorama | Normalized gene expression matrix | Scanorama version 1.6 [7] |
| ComBat | Normalized gene expression matrix | sva version 3.38.0 [13] |
| Harmony | Normalized feature reduction vectors | Harmony version 1.0 [8] |
| Liger | Normalized feature reduction vectors | Liger version 0.5.0 [9] |
| ComBat-seq | Normalized gene expression matrix | sva version 3.38.0 [12] |
| scMerge | Normalized gene expression matrix | scMerge version 1.6.0 [14] |
| Canek | Normalized gene expression matrix | Canek version 0.1.7 |

**Table 2.** Cell type distribution on simulated data.

| Method | Path cells |  | Group cells |  |  |
| --- | --- | --- | --- | --- | --- |
|  | Cell-1 | Cell-2 | Cell-3 | Cell-4 | Cell-5 |
| Batch-1 | ✓ | ✓ | ✓ | - | - |
| Batch-2 | - | ✓ | - | ✓ | - |
| Batch-3 | ✓ | - | - | - | ✓ |

### Application to real datasets

Next, we compared Canek with other methods on 3 real datasets: Tabula Muris spleen, human pancreatic islets, and interferon beta stimulation [17, 19–23].

First, we tested a scenario in which the same sample was used with two different technologies simultaneously. For this we integrated the droplet and FACS batches from the Tabula Muris spleen datasets [17]. Supplementary Figure 6 shows UMAP plots for the Uncorrected data, and the correction done by Canek and the other 8 correction methods. Except for scMerge, which merged some cell populations, all the methods successfully integrated the datasets, with cells from the same type found in the same clusters. This demonstrates that Canek can integrate datasets even from different technologies.

Next, we tested the scenario in which similar tissues were used with different technologies. For this we integrated eight human pancreatic islet datasets from five different technologies. Figure 5a shows the Uncorrected data, where the batch effect caused the cells to cluster by batch. The results for all methods are shown in Supplementary Figure 7. Figure 5 highlights the results from Canek and Seurat. Canek (Figure 5b) was able to integrate the batches, but some differences remained. Other methods like Seurat (Figure 5c) and MNN (Supplementary Figure 7g) mixed the batches almost perfectly. Interestingly, the differences remaining in Canek integration are correlated with disease state (Supplementary Figure 8), with some of the samples containing type 2 diabetes whereas other containing only healthy individuals. Therefore, we tentatively speculate that the observed differences may, indeed, be due to true biological differences.

**Figure 5.**
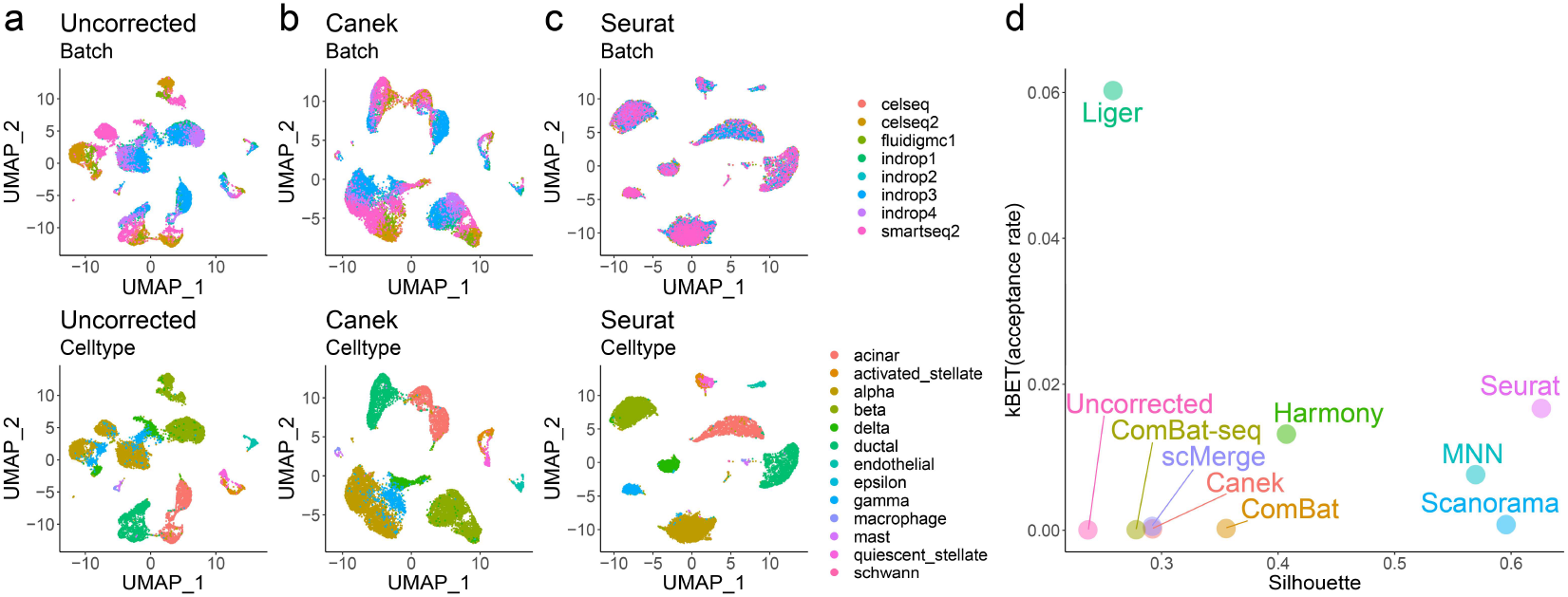
Integration datasets from different technologies. Eight pancreatic datasets obtained using different technologies were corrected. **a)** Batch effects caused the cells to cluster by batch instead of by cell type. **c-f)** The batches were integrated using different methods.

Finally, we evaluated the scenario wherein two samples from different conditions were assayed using the same technology. For this we integrated a dataset obtained from PBMCs with and without interferon-beta stimulation [24]. In this scenario, differences between the same cell types due to the stimulation were expected. Supplementary Figure 9a shows that in the Uncorrected data, cells separate by batch. Supplementary Figures 4b-j shows the correction done by Canek and 8 other methods. After integrating with Canek, B cells and T cells were almost completely integrated but some differences remained. Differences in monocytes and dendritic cells in the stimulated vs. non-stimulated cells were more prominent. This is in agreement with experiments showing that interferon beta induces stronger changes in gene expression in monocytes compared to T cells [25].

### Integration of a human lung dataset

To evaluate how Canek performs on the task of integrating samples from replicated experiments, we used a human lung single cell dataset with 78 samples including IPF (n = 32; idiopathic pulmonary fibrosis), COPD (n = 18; chronic obstructive pulmonary disease), and control donors (n = 28) [26]. This dataset consisted of 312,928 cells distributed over 107 sequencing libraries that we treated as different batches. Figure 6a–b shows that Canek integration resulted in good mixing among libraries while preserving the cell populations identified in the original publication. These cell types closely correlated with cell clusters based on Canek integration (Figure 6c). Most cells were distributed evenly among all cell types and disease conditions (Figure 6d). Enrichment and depletion in cell populations associated with disease were preserved (Supplementary Figure 10). A group of macrophages enriched in IPF in clusters 12 (interstitial macrophages expressing matrix metallopeptidase 9; MMP9, Figure 7e–g) and 16 (alveolar macrophages) showed cells in a transitional state almost exclusively from IPF donors [26]. This demonstrates that Canek can integrate a high number of replicated datasets while preserving biologically meaningful information.

**Figure 6.**
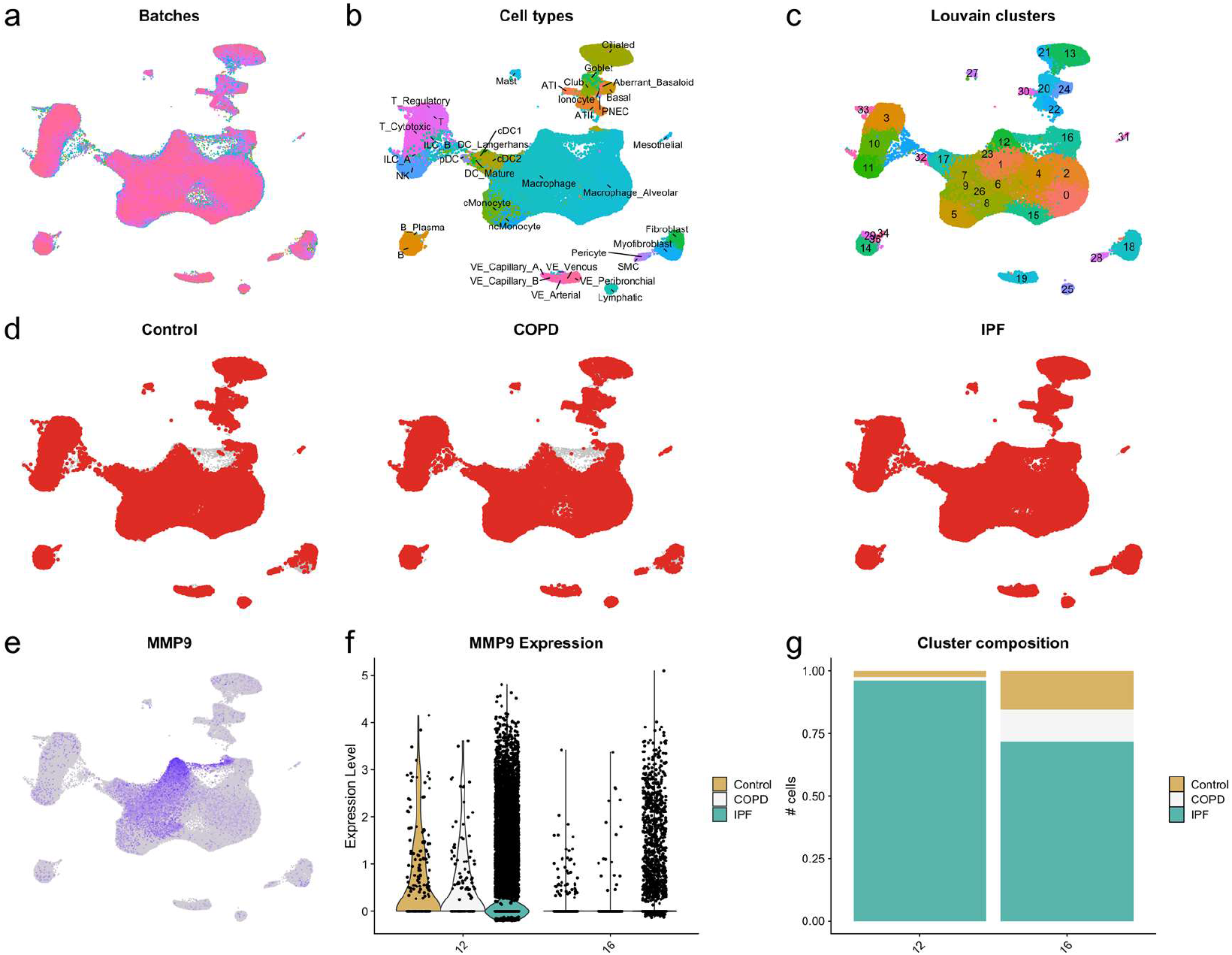
Integration of the lung dataset from Adams (2020). a) UMAP plot showing the mixing between batches. b) Cell populations described in the original publication are preserved. c) Clustering based on Canek integration matches with cell populations. d) Distribution of cells by disease condition shows even distribution except for a group of IPF specific cells in clusters 12 and 16. e) MMP9 is highly expressed in cluster 12 (interstitial macrophages), f) in IPF donors. g) Cluster 12 and 16 are enriched in cells from IPF.

**Figure 7.**
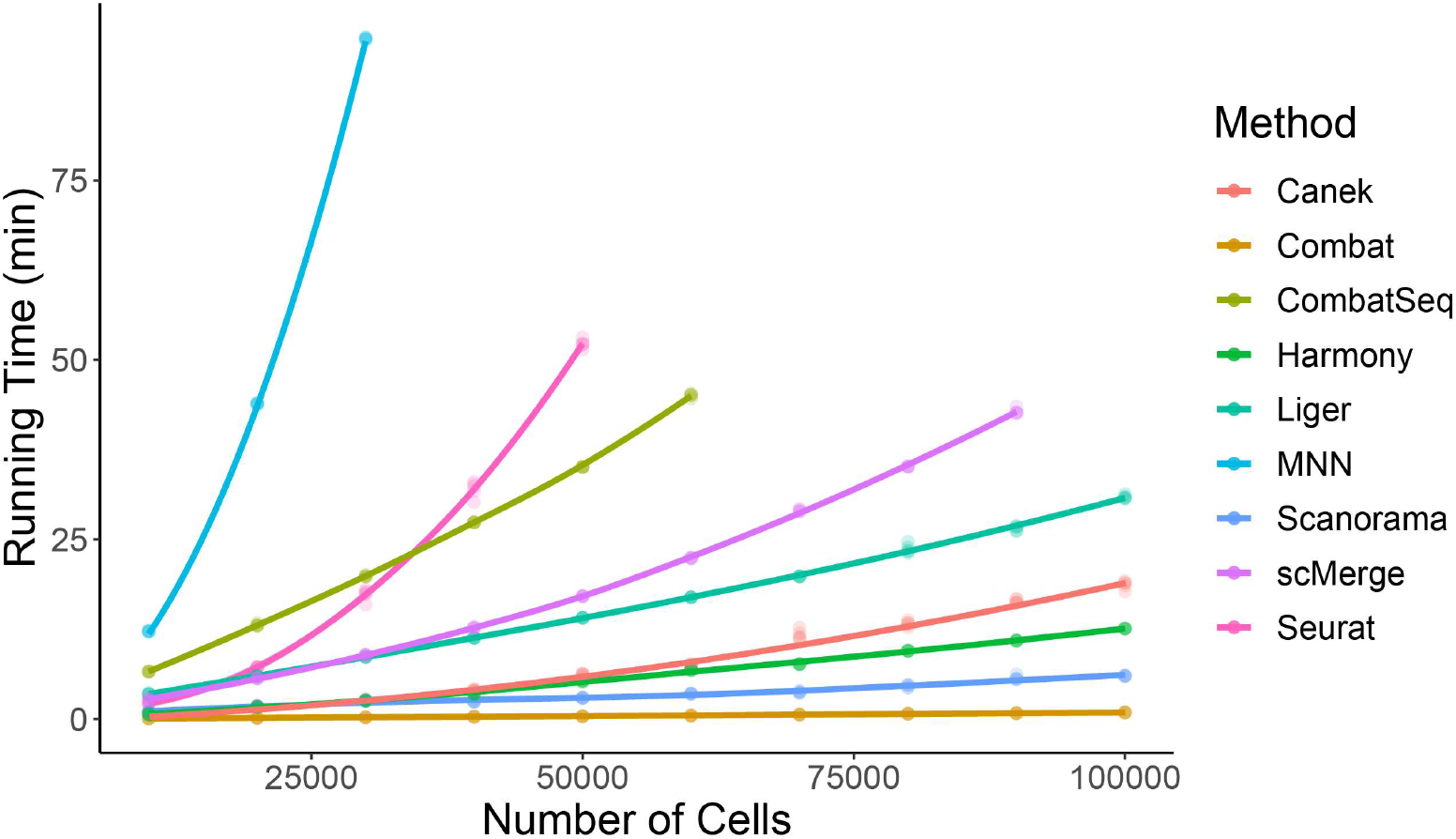
Runtime benchmark of Canek and other eight batch correction methods. Each method was run 5 times on different datasets with the number of genes fixed to 2k and the number of cells varying in a range of 5k to 100k. The color code differentiates each of the methods, the dots represent the runtime, and the lines represent the time increasing trends. Canek displayed a linear time increase over these conditions.

### Computational performance

To compare the computational performance and scalability of Canek and eight other batch correction methods, we simulated datasets using splatter and recorded the integration time. We fixed the number of genes to 2,000 and varied the number of cells from 10k to 100k. Figure 7 shows run times as a function of the number of cells. The fastest method was Combat, followed by Scanorama, Harmony, and Canek, all of which showed near linear run time dependence and ability to integrate 100k cells in under 20 min. On the other side of the spectrum MNN, Seurat, and ComBat-seq showed a strong dependence of run time on data size. These results demonstrated that Canek is a scalable method that can integrate hundreds of thousands of single-cell transcriptomes efficiently.

## Discussion

Existing batch effect correction methods focus on integrating single-cell transcriptomics datasets obtained from different technologies and/or species, minimizing the differences among batches to obtain correlated cell types. While these frameworks offer a powerful solution to integrate datasets with strong differences between batches, they may also introduce significant biases due to over-correction. This represents a potential problem when these methods are applied to datasets where we wish to preserve biological differences (e.g., replicated experiments obtained with the same technology). Over-correction could negatively affect downstream tasks such as clustering or differential gene expression analysis. Canek provides an unbiased batch effect correction method for single-cell transcriptomics data that is suited for such integration of experimental replicates. We focused on preserving the inherent biological structure while being flexible enough to deal with small non-linear differences that might appear on heterogeneous datasets. We applied Canek to simulated and real datasets and showed its ability to correct batch effects without masking real biological signals. We also tested Canek on a pseudo-batch scenario with no batch effect and observed that it preserved the biological structure and introduced the least undesirable bias among tested methods.

We further showed that Canek successfully integrated datasets from different technologies (e.g., the Tabula Muris spleen dataset). Depending on the nature of the dataset, Canek did not necessarily lead to the best batch mixing (e.g., in the human pancreatic islet integration). However, latent variables other than batch effects (i.e., disease condition) may influence the integration of these datasets. It is an open question how to integrate complex datasets with cofounding variables.

The main goal of Canek is to enable efficient and unbiased integration of replicated experiments. Thus, we applied Canek to a large dataset from human lung disease. We identified enrichment of cell populations reported in the original publication, including cells in an apparently transitional state between interstitial to alveolar macrophages. This showed that Canek was able to integrate large numbers of replicated experiments while preserving biological information.

As single-cell RNA-seq from replicated experiments using the same technology become more common, batch effect correction methods that conserve local differences will become more important. Canek provides a solution to this problem with an unbiased and computationally efficient batch effect correction.

## Methods

### Canek workflow

Figure 2 shows the workflow for correcting a pair of batches. We define one of the batches as the *query batch* and the other one as the *reference batch*. We correct the cells from the query batch to match the cells from the reference batch. When correcting more than two batches we perform an optional hierarchical optimization of batch order (see *Hierarchical integration* section). The main steps of Canek’s workflow are:

1. Obtaining **batch effect observations** using mutual nearest neighbors (MNN) pairs.
2. **Clustering** the query batch to define local groups of cells.
3. Calculating a batch effect **correction vector** for each cluster.
4. Obtaining a **fuzzy correction** by smoothing the transitions between the local correction vectors.

Canek expects input datasets to be log normalized. The output dataset retains the same dimensionality (number of genes) as the input batches.

#### Batch effect Observations

The first step is to identify what we call batch effect observations. This is the gene expression differences of a set of cells from the reference and query batches that will enable us to estimate the batch effect.

To speed up computation we calculate the first 50 principal components (PCs) [27] using the *prcom_irlba* function from the ilrba R package [28]. This lower dimensional space is used to identify MNNs, and in the clustering and fuzzy correction steps. However, during the calculation of the correction vector step, we use the original input datasets.

We calculate mutual nearest neighbors (MNN) pairs [4] using 50 PCs to obtain batch effect observations. We assume that at least one cell is shared between the batches to integrate. The MNN pairs are defined by the intersection of the *crossed k nearest neighbors* for each cell of two input batches. For example, for a cell *c*_1_ from batch one, we find the k closest cells from batch two, and for cell *c*_2_ from batch two, we find the k closest cells from batch one. If *c*_1_ and *c*_2_ are mutually contained on each other’s nearest neighbor set, they are considered as a MNN pair. In Canek, to identify MNN pairs we first find the crossed 30 nearest neighbors of the query and reference batches using the *get.knn* function from the FNN R package [29]. We then select those cells that fulfill the MNN criteria to form cell pairs. We treat the gene expression differences from these pairs as observations of the batch effect.

#### Clustering

Following Haghverdi et al. (2018) [4] we assume that the batch effect is almost orthogonal to the biological space, and that the variations due to the batch effect are smaller than the biological variation (see Supplementary material of [4] for a deeper discussion of these assumptions). Small variations to this orthogonality assumption can be caused by noise or by non-linearities. A common way to deal with non-linear dynamics is to linearize over bounded regions [30], to solve each of these local problems, and, if necessary, to join all the pieces back into a non-linear global solution. Following this idea, we partition the query batch into clusters, which we define as a bounded set of related cells, using the Louvain algorithm implemented in the igraph R package [31]. By default, clustering is done using the first 10 PCs.

#### Correction Vector

Following our *local orthogonal batch effect* assumption, *for each* cluster we state the relation:

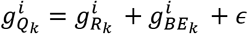

where *g*^*i*^, *i* = 1,…, *n*, is the log-normalized gene expression level of the *n* genes from the input batches. The batch effect 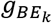 is represented as an additive value in the query batch 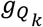 in terms of the same gene in the reference batch 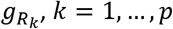, being *p* the number of MNN pairs from the membership under analysis. Finally, *ϵ* represents a normally distributed random error term with mean zero and standard deviation *σ*, which we assume to be independent of *g*^*i*^ on each cluster. Thus, using

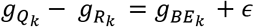

on each gene *i*, the term 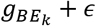 would be normally distributed with mean *μ* = *g*_*BE*_ and standard deviation *σ*. Accordingly, a good estimation of the batch effect would be the mean of the gene expression subtraction between MNN cells pairs (e.g. 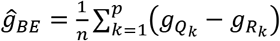). But there is a complication with this approach, since *erroneous pairs* between cells from distinct but related cell types could be formed, resulting in the incorrect integration of dissimilar subpopulations [5]. To tackle this problem, reasoning that abnormal pairs would appear as outliers to the normal distribution of 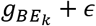, we estimate a correction vector

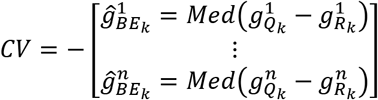

where the function *Med* represents the statistical median, which is less affected by outliers than the mean. Canek uses this approach by default to reduce the impact from outliers, but it is possible to perform an optional filtering step (with extra computational cost) based on the interquartile range to detect MNN outliers (see *Filtering* section).

#### Fuzzy correction

From the steps described above, each cell from the same cluster will be assigned the same correction vector. We use fuzzy logic to smoothly join the cluster-specific corrections into a cell-specific one, where each cell has a unique correction vector (see Supplementary Figure 12). Within this fuzzy logic framework, the clusters previously identified will be considered as memberships.

Using the PCs of the query batch, we create a minimum spanning tree (MST) over the memberships’ center points (*MC* s) using the *mst* function from the R package igraph [31] (Supplementary Figure 11a,b). For each edge of the MST, we construct a pair of membership functions (*MF*s). These *MF*s are used to calculate a fuzzy score for the cells (Supplementary Figure 11c,d). For example, let us consider an edge that joins the centers of memberships number 1 (*MC*_1_) and 2 (*MC*_2_). For each cell *j* that belongs to memberships 1 or 2, we define the vector *V*_*j*_ as a vector for cell *j* from *MC*_1_ in the PCs embeddings. Similarly, let 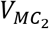 be the vector corresponding to *MC*_2_. Then, we obtain the scalar projection *p*_*j*_ for each cell *j* onto the line connecting *MC*_1_ and *MC*_2_ as:

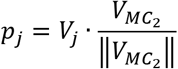

where the operator . denots the dot product, and 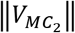 is the length of 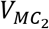. We then construct the *MF*s (i.e., *MF*_1_ and *MF*_2_) as

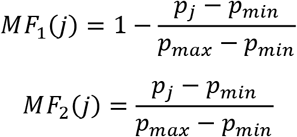

Here, *p*_*max*_ and *p*_*min*_ are the maximum and the minimum of the scalar projections of the cells in the memberships (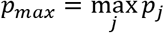 and 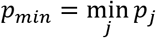). In this way, the membership function *MF*_1_ (*MF*_2_) takes the maximum value 1 (the minimum value 0) for *p*_*min*_ and the minimum value 0 (the maximum value 1) for *p*_*max*_, respectively, and linearly interpolates for the other values of the projections. (Supplementary Figure 12).

We calculate cell specific correction vectors *CV*_*j*_ by using the Takagi-Sugeno approach [32] to combine the membership’s correction vector *CV*^*(l)*^ (see *Correction Vector* section) with the membership functions:

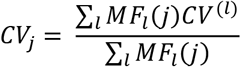

When a membership *l* is connected to several edges, we use the average of the membership functions *MF*_*l*_ defined for all the edges associated with membership *l*.

Even though the fuzzy scheme is applied in a low dimensional representation, the final output is in the original dimensionality of the input datasets. We recommend using Canek with the fuzzy step, but to skip it users can set the boolean parameter *fuzzy to FALSE*. In this case the final integration will be done using a membership-specific correction instead of a cell-specific one.

### Hierarchical integration

We define a hierarchical integration when Canek is applied to more than two input batches. We first sort the batches by cell number in descending order and use the batch with the higher number of cells as the first reference batch. To determine the query batch, we prioritize to integrate first related batches as they would have a higher number of MNN pairs. The query batch is therefore chosen as the batch sharing the highest number of MNN pairs with the reference. For this, we obtain their first three PCs using the prcomp_irlba function in the irlba R package [28], find the MNN pairs and select the query batch as the one with the highest number of pairs. Once the reference and the selected query batch are integrated, we define the integrated batch as the new reference, and again select the query batch following the same procedure as before. We continue this process until all the input batches are integrated. The hierarchical integration is optional and can be deactivated by setting the boolean parameter *hierarchical* to FALSE. In this case, the order of integration follows the order in the input list.

### Filtering

We assume that erroneous MNN pairs would appear as outliers from the normal distribution 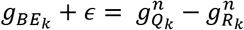 of (see *Correction Vector* section). We use the median function to reduce the impact of these outliers on the correction vector estimation. In addition, the user can select an extra filtering step based on the interquartile range:

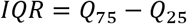

where *Q*_75_ and *Q*_25_ are the 75^th^ and 25^th^ percentiles of the distribution of the *p* MNN pairs’ Euclidean distance *d(k), k* = 1,…, *p*. Therefore, we will select and filter any outlier MNN pairs as:

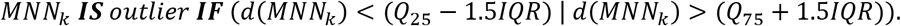

### Analysis details

#### Data pre-processing

We performed the same data pre-processing for all the analyses. We implemented the “Standard Workflow” from the Seurat R package [33], which involves:

- **Normalization**: using the function NormalizeData. Gene expression levels are divided by the total number of transcripts and multiplied by 10,000. The results are then log normalized.
- **Identification of high variable features**: using the function FindVariableFeatures. Genes that show high variations among cells are selected using the vst method.

#### Batch-correction algorithms

Table 1 lists the batch correction methods used.

To objectively compare the batch effect correction methods, we used the same pre-processed data and the same variable genes on each method. We obtained the variable genes from Seurat’s integration using the *VariableFeatures(assay=“integrated”)* function from the Seurat R package [33] and used them to subset the pre-processed uncorrected datasets. We implemented the integration methods as follows:

##### Seurat

We used the *FindIntegrationAnchors* and *IntegrateData* functions with default parameters from the Seurat R package [33].

##### Canek

We used the *RunCanek* function from the Canek R package with default parameters.

##### MNN

We used the *mnnCorrect(cos.norm.out = FALSE)* function with default parameters from the batchelor Bioconductor package [4].

##### Scanorama

We used the *scanorama.correct(return_dense=TRUE)* with default parameters from the scanorama Python library [7].

##### ComBat

We used the *ComBat* function with default parameters from the sva R package [13].

##### Harmony

We used the *RunHarmony* function with default parameters from the harmony R package [8].

##### Liger

We used the *RunOptimizeALS(k=20, lambda=5)* and the *RunQuantileNorm* functions with default parameters from the SeuratWrappers R package [9].

##### ComBat-seq

We used the *ComBat_seq* function with default parameters from the sva R package [12].

##### scMerge

We used the *scMerge* function with default parameters from the scMerge R package [14]. For the parameter *kmeansK,* we used the number of cell types when known, otherwise the number of clusters.

After correcting batch effects, we scaled each of the integrated and uncorrected datasets using the *ScaleData* function from the Seurat R package, except for Harmony and Liger integrations, as their output is already a low dimensional space.

#### Dimensionality reduction

We obtained the principal components [27] from the corrected and uncorrected datasets using the RunPCA function from the Seurat R package. We used the first 10 PCs as the standard in all the tests. In the case of the UMAP representation [15], we applied the *RunUMAP* from the Seurat R package to the selected PCs.

#### Simulated data

We used the *splatSimulate* function from the splatter Bioconductor package [18] to simulate three batches with batch effect. Splatter allows us to simulate cell types whether as groups or paths. Because we wanted to assess cell type preservation on a mixed population scenario with clearly defined groups along with a differentiation process, we simulated paths and groups separately and merged them. Then, we manually removed cells such that the batches shared only one cell type. The final cell type composition is:

After removing the cell types, the number of cells per batch is: 1,671 cells for Batch 1, 975 cells for Batch 2 and 964 cells for Batch 3, all of them with the same 2,000 genes.

To obtain the gold standard without batch effects, we used *splatSimulate(batch.rmEffect = TRUE)* and removed the same cells as the simulations with batch effects.

### Running time benchmark

We used the *splatSimulate* function from the *Splatter* Bioconductor package [18] to simulate two datasets with a 2,000 genes and a varying number of cells in the range of 10k to 100k. Each dataset contained three cell types with appearance probabilities of 0.3, 0.3, and 0.4 respectively. We applied each of the correction methods five times on each of the simulated datasets and recorded the time. We used the *geom_smooth* function from the ggplot2 R package [34] to plot the time trend lines.

#### Public datasets

Table 3 lists the public datasets we used.

**Table 3.**
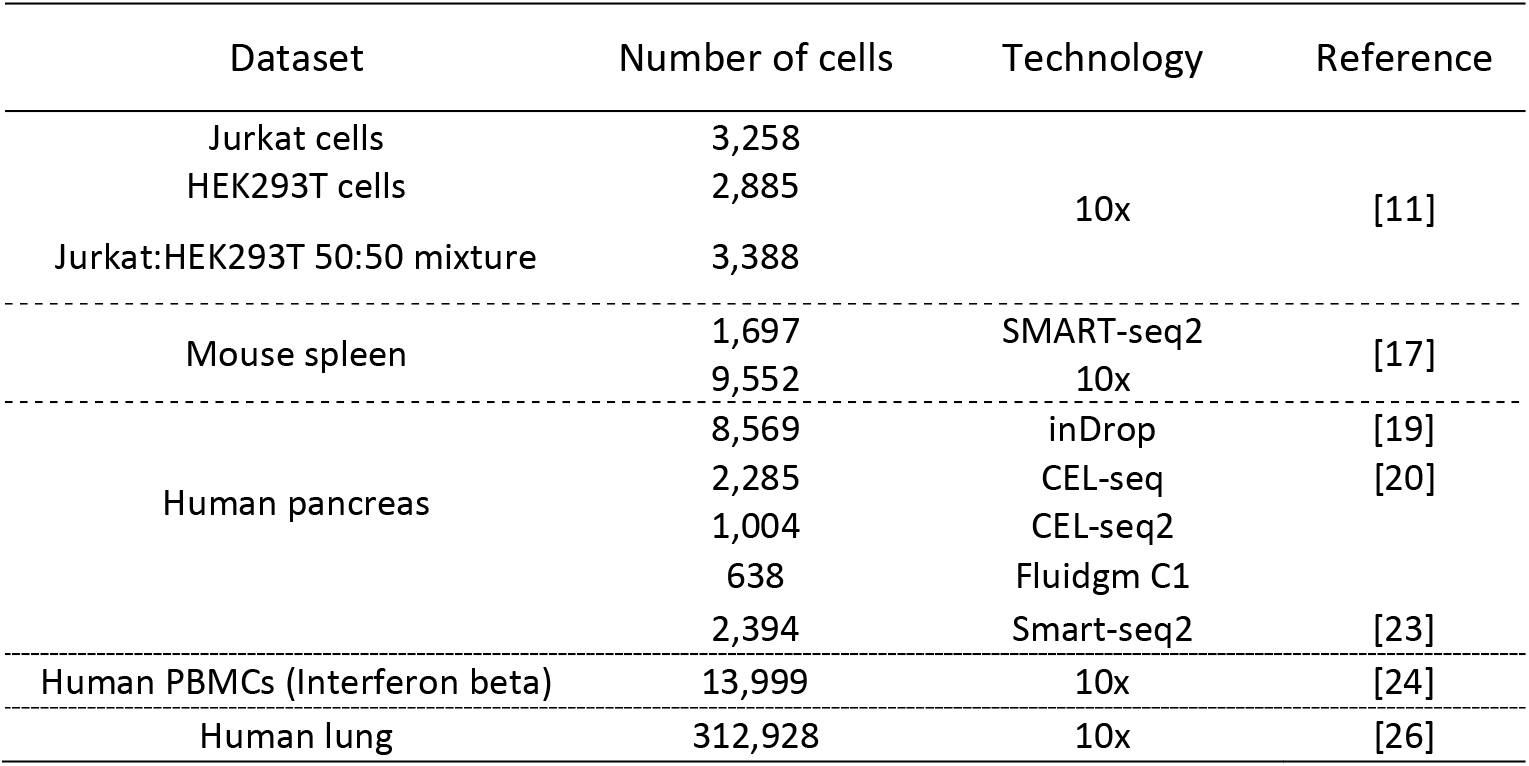
Public datasets used.

### Jurkat/t293 data analysis

We used the following publicly available datasets:

- 293T cells. https://support.10xgenomics.com/single-cell-gene-expression/datasets/1.1.0/293t_3t3
- Jurkat cells. https://support.10xgenomics.com/single-cell-gene-expression/datasets/1.1.0/jurkat
- 50:50 Jurkat:293T cell mixture. https://support.10xgenomics.com/single-cell-gene-expression/datasets/1.1.0/jurkat:293t_50:50

In the 50:50 Jurkat:293T cell dataset, we used the *kmeans* function from the *stats* R package [35] with k = 2, checked the expression of the XIST and CD3D3 genes and assigned the appropriate cell type labels.

### Spleen data analysis (pseudo-batch)

- We use the publicly available dataset from Tabula Muris [17] https://ndownloader.figshare.com/files/13090478.

### Pancreatic data analysis

- We obtained the five public datasets [19–23] from the SeuratData R package[5] and used the provided cell type labels.

### PBMC unstimulated and IFN-β-stimulated data analysis

- We obtained the public datasets [24] from the SeuratData R package [5] and used the provided cell type labels.

### Human lung dataset

Single cell RNA-seq data and associated metadata from human lung was downloaded from GSE136831, loaded into R and converted into a Seurat object [5]. This object was converted into a list of objects with the SplitObject function using Library_Identity as batch variable. The standard processing workflow was used to normalize and find highly variable features in each of the libraries (see Data preprocessing above). This data was passed to the function RunCanek with hierarchical integration set to FALSE. The integrated dataset was scaled using ScaleData and Principal Component Analysis dimensions were calculated with RunPCA. The top 25 principal components were used to calculate UMAP using RunUMAP, and to create the Shared Nearest Neighbors graph with FindNeighbors. Clustering was done using FindClusters with the Louvain algorithm and a resolution of 1. Cells annotated as multiplets in the original publication were removed.

#### Metrics

We evaluated the results from the batch-correction methods by scoring the mixing between batches with the k-nearest-neighbor batch effect test (kBET) [36], and the cell purity preservation with the average Silhouette width (Silhouette) [37]. We used the kBET and batch_sil functions from the kBET R package [36].

The kBET metric provides a rejection rate within 0 and 1 after testing batch mixing at the local level. The kBET’s score could be affected by the choice in the number of k-nearest neighbors (kNN). To objectively assess the different integration methods, following the idea of Tran et al. [3], we obtained the mean cell number of the datasets and performed the scoring by fixing the kNN size as the 5%, 15%, and 30% of this mean. To ease the interpretation of this metric, we calculated an “acceptance rate” by subtracting the rejection rate from 1.

We used the Silhouette coefficient to assess cell purity after integration [37]. This metric analyzes the separation among cells from the same cluster as compared with cells from other clusters. Let *a*(*i*) be the average Euclidean distance of cell *i* to all other cells from the same cluster as *s*(*i*), then the Silhouette width *s*(*i*) is defined as:

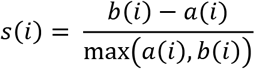

where *b*(*i*) is calculated as

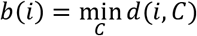

being *d*(*i, C*) the average distance of cell *i* to all the other cells assigned to different clusters *C*. A higher score means a longer separation between clusters and a lower score means a shorter separation. We used the cell type labels provided for each dataset as inputs to the Silhouette coefficient, except on the pseudo-batch experiment, where we obtained the cluster labels using the *FindNeighbors* and *FindClusters(resolution = 0.5)* functions from the Seurat R package.

For both kBET and Silhouette metrics we used the “harmony” and “iNMF” embeddings from Harmony and Liger corrections respectively. For the rest of the methods and the Uncorrected data we used the first 10 PCs.

## Data availability

The datasets used for the simulation and pseudo-batch tests are available from figshare: https://figshare.com/projects/Canek/122638.

## Code availability

Canek is implemented as an R package and is available from GitHub: https://github.com/MartinLoza/Canek. Code used to replicate the analyses presented in this paper is available from: https://github.com/MartinLoza/WorkflowsCanek

## Acknowledgements

We thank Dr. Alexis Vandenbon for valuable comments on our manuscript.

## Author contributions

ML conceived this work with contributions from ST and DD. DMS provided guidance for benchmarking and performance evaluation. ML and DD implemented the methods. ML performed the analyses with contributions from DD. ML and DD wrote the manuscript with contributions from ST and DMS. All authors edited and approved the submitted manuscript.

