## Supplementary figures for "Unbiased integration of single cell transcriptome replicates"

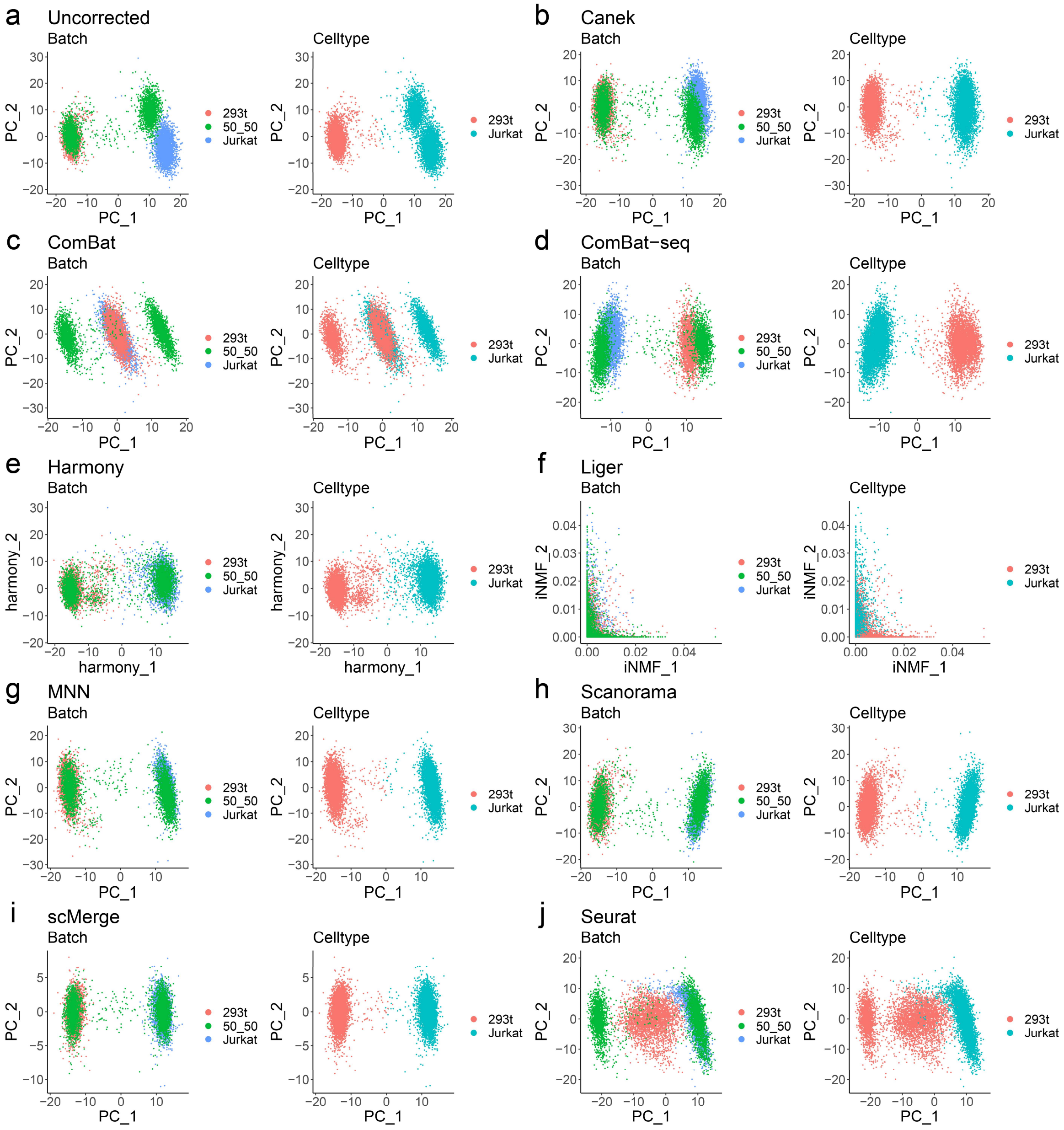


*Supplementary Figure S1. PCA plots for batch and celltype of the Jurkat/293t cells mixture dataset a) before and b-j) after batch correction.*

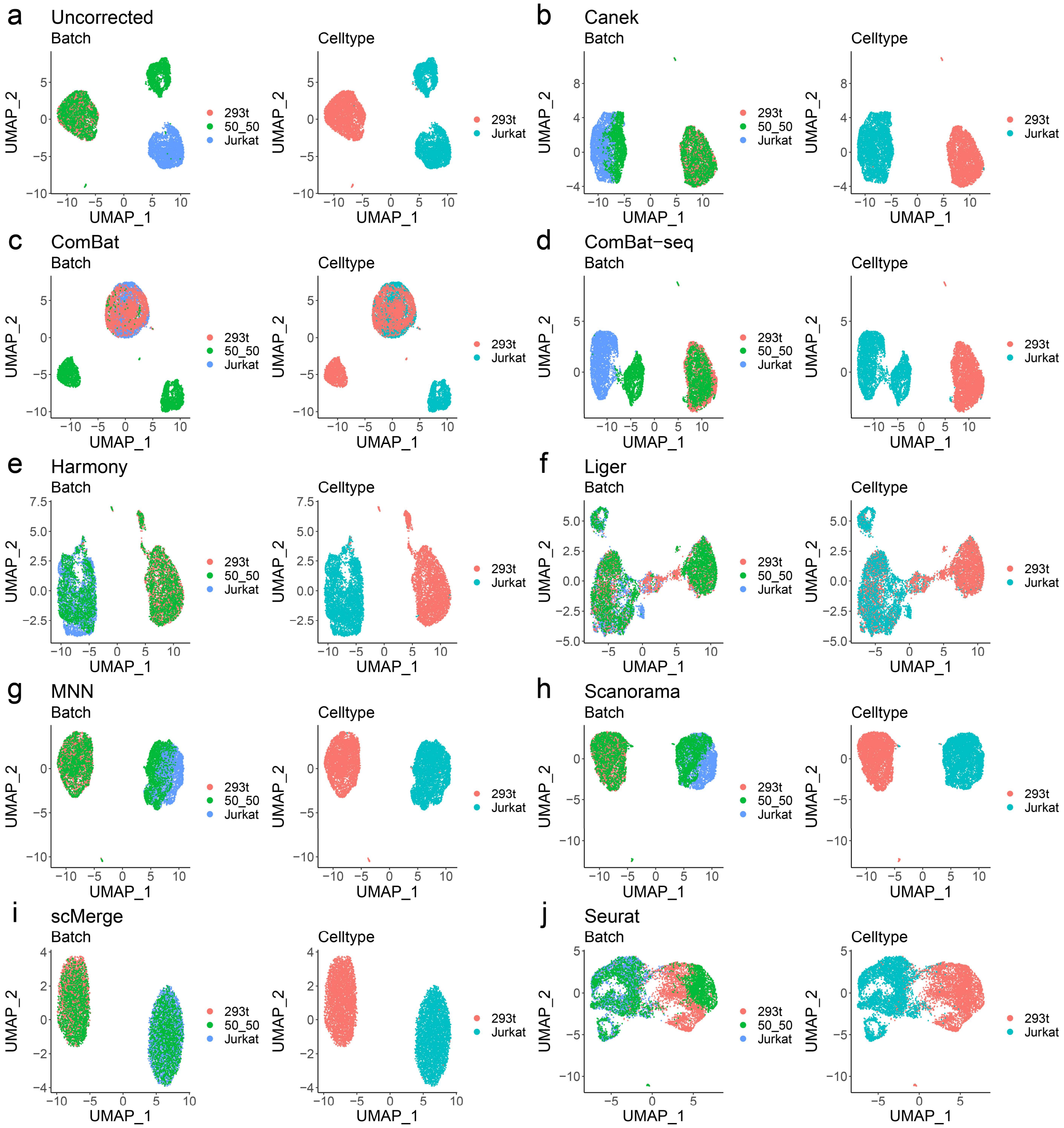


*Supplementary Figure 2. UMAP plots for batch and celltype of the Jurkat/293t cells mixture dataset a) before and b-j) after batch correction.*

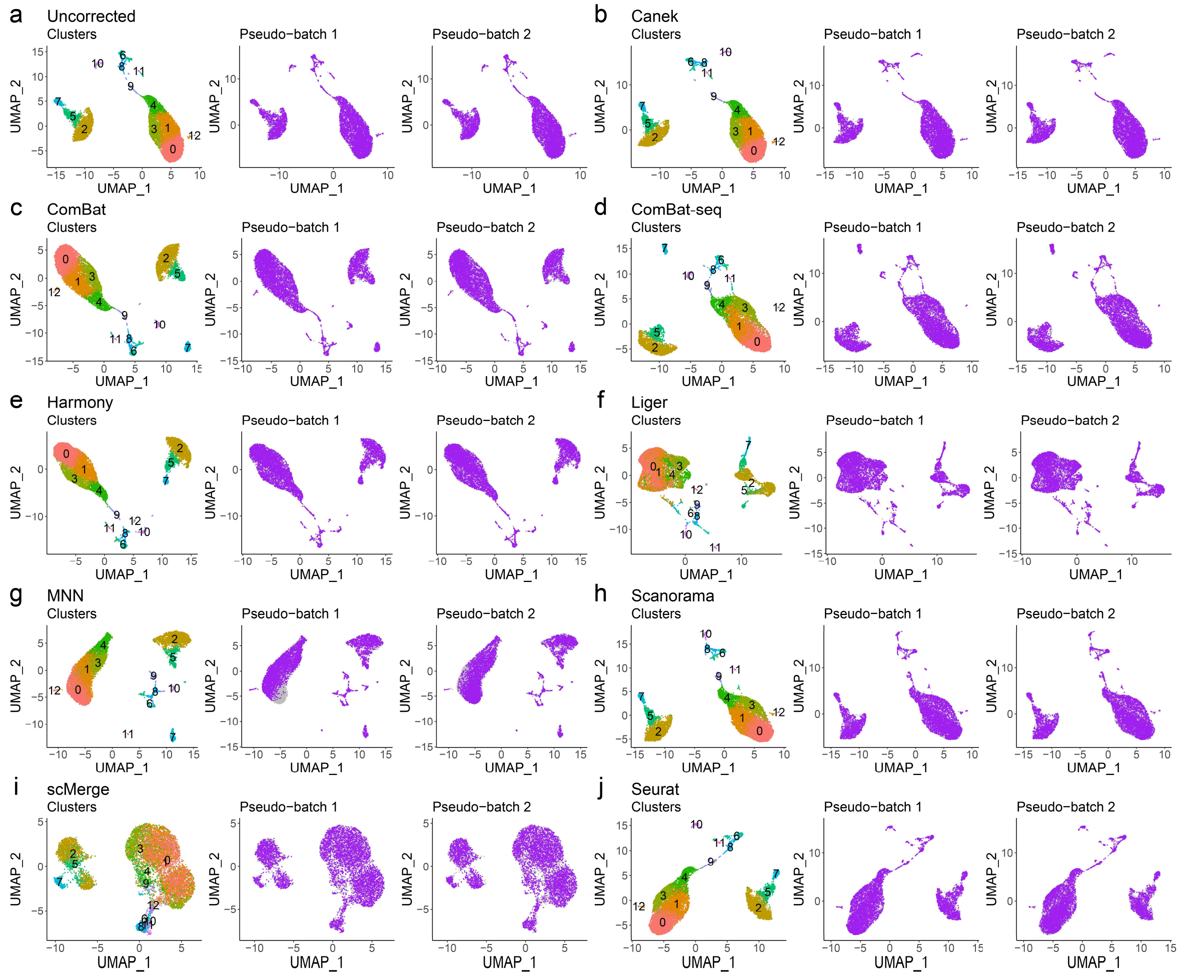


*Supplementary Figure 3. UMAP plots for all methods in the pseudo-batch experiment.*

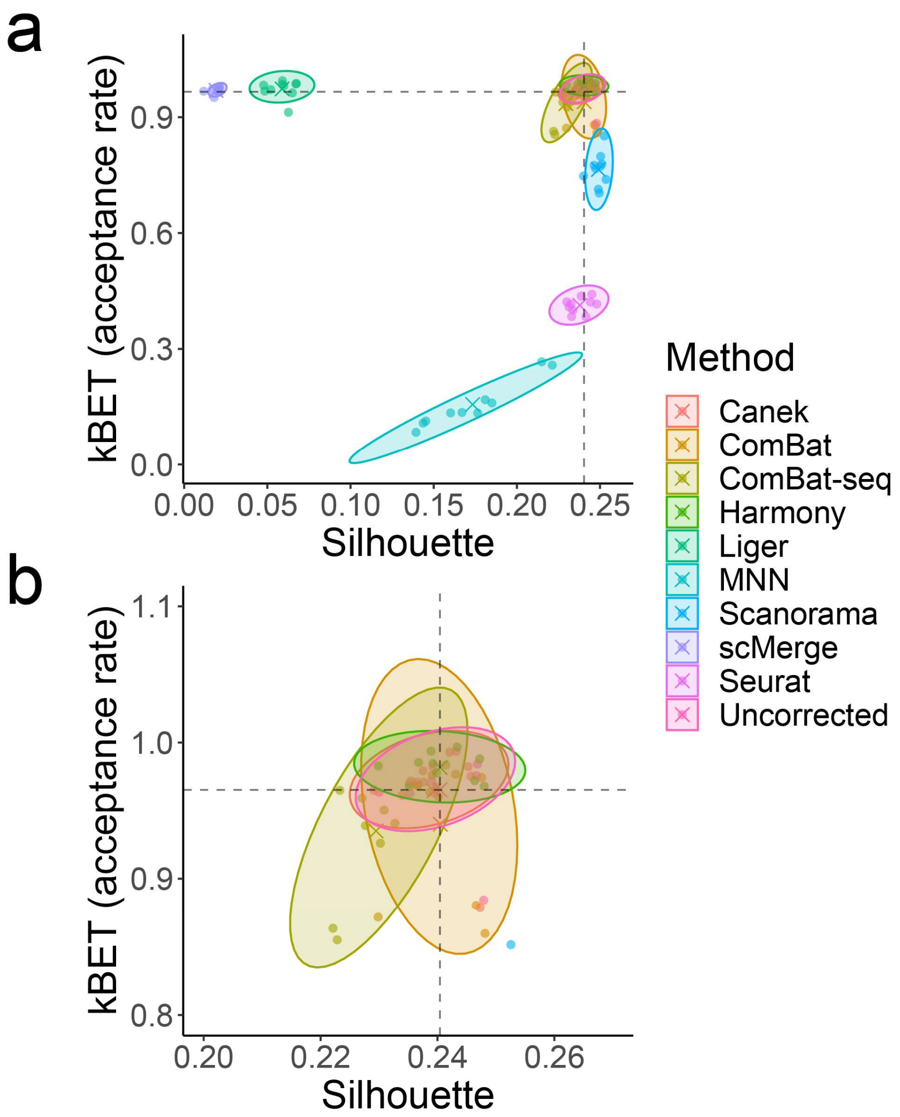


*Supplementary Figure 4. **a)** Silhouette vs. kBET scores for the 10 pseudo-batch experiments together with the average of each kBET/silhouette replicate. **b)** Zoom in to the methods closer to the metrics for the Uncorrected dataset (intersection of dashed lines).*

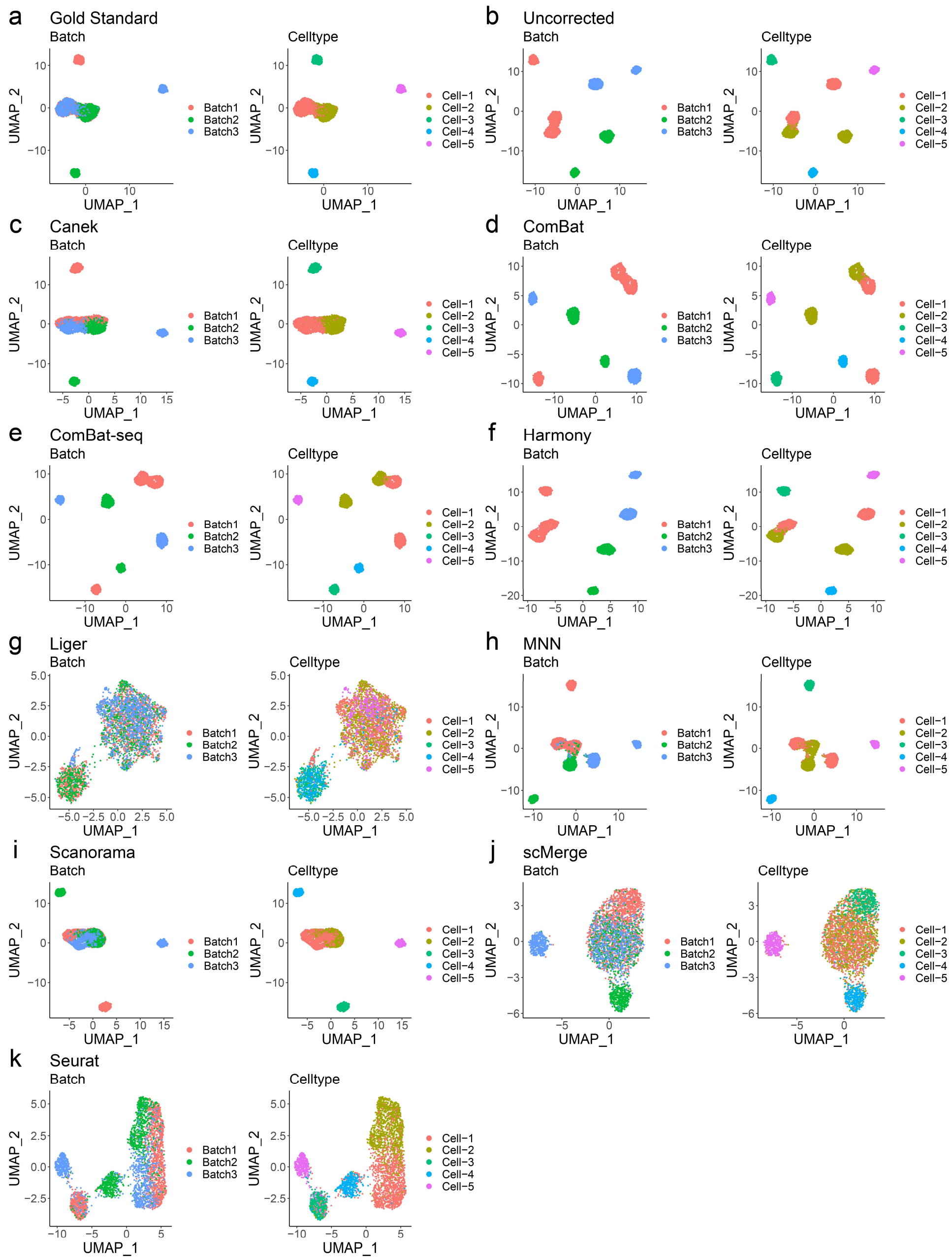


*Supplementary Figure 5. UMAP plots for the Gold Standard, Uncorrected dataset, and the result of applying batch correction methods. The left-side plot shows the batches while the right-side plot shows the cell types.*

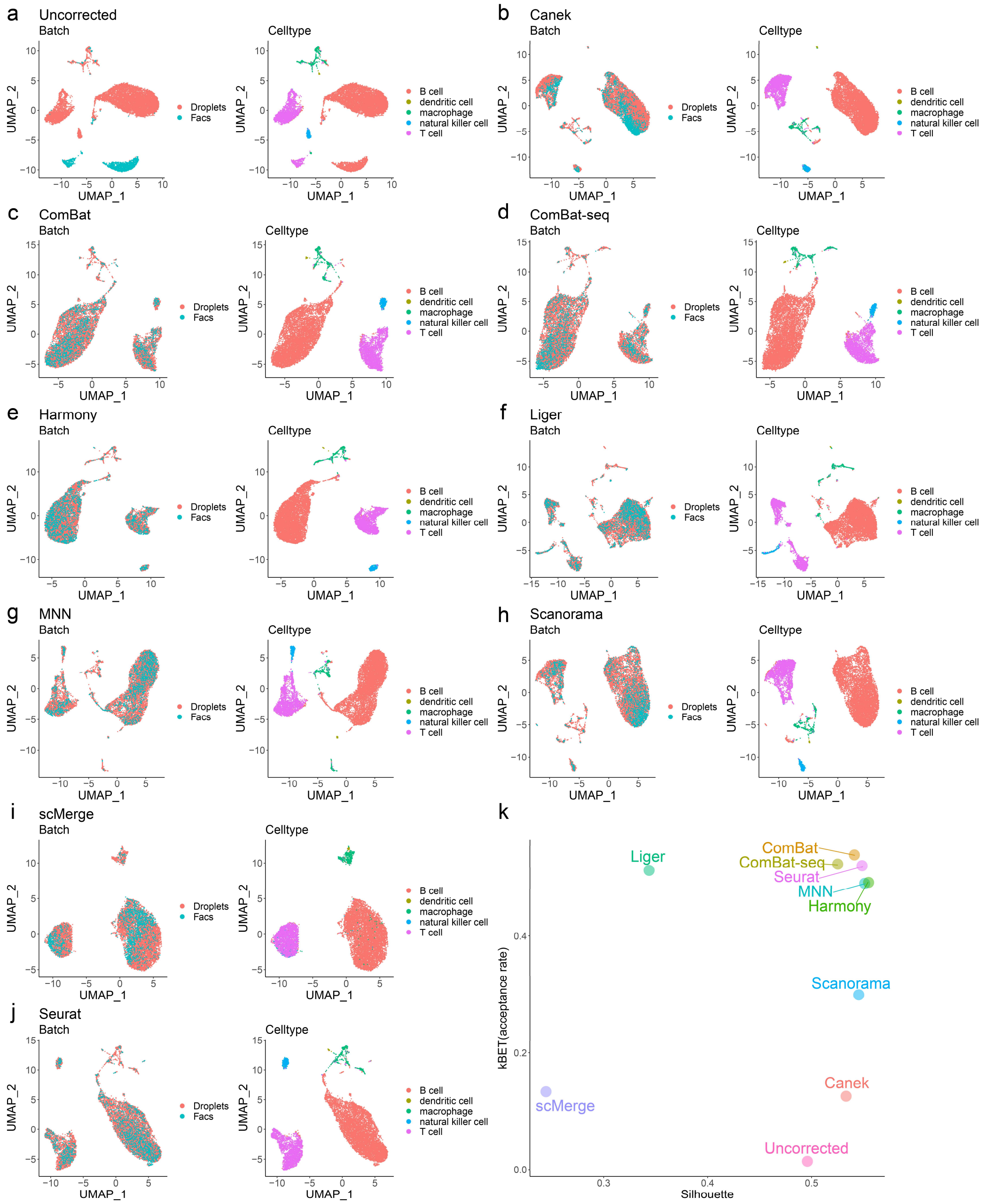


*Supplementary Figure 6. Integration of Tabula Muris spleen datasets.*

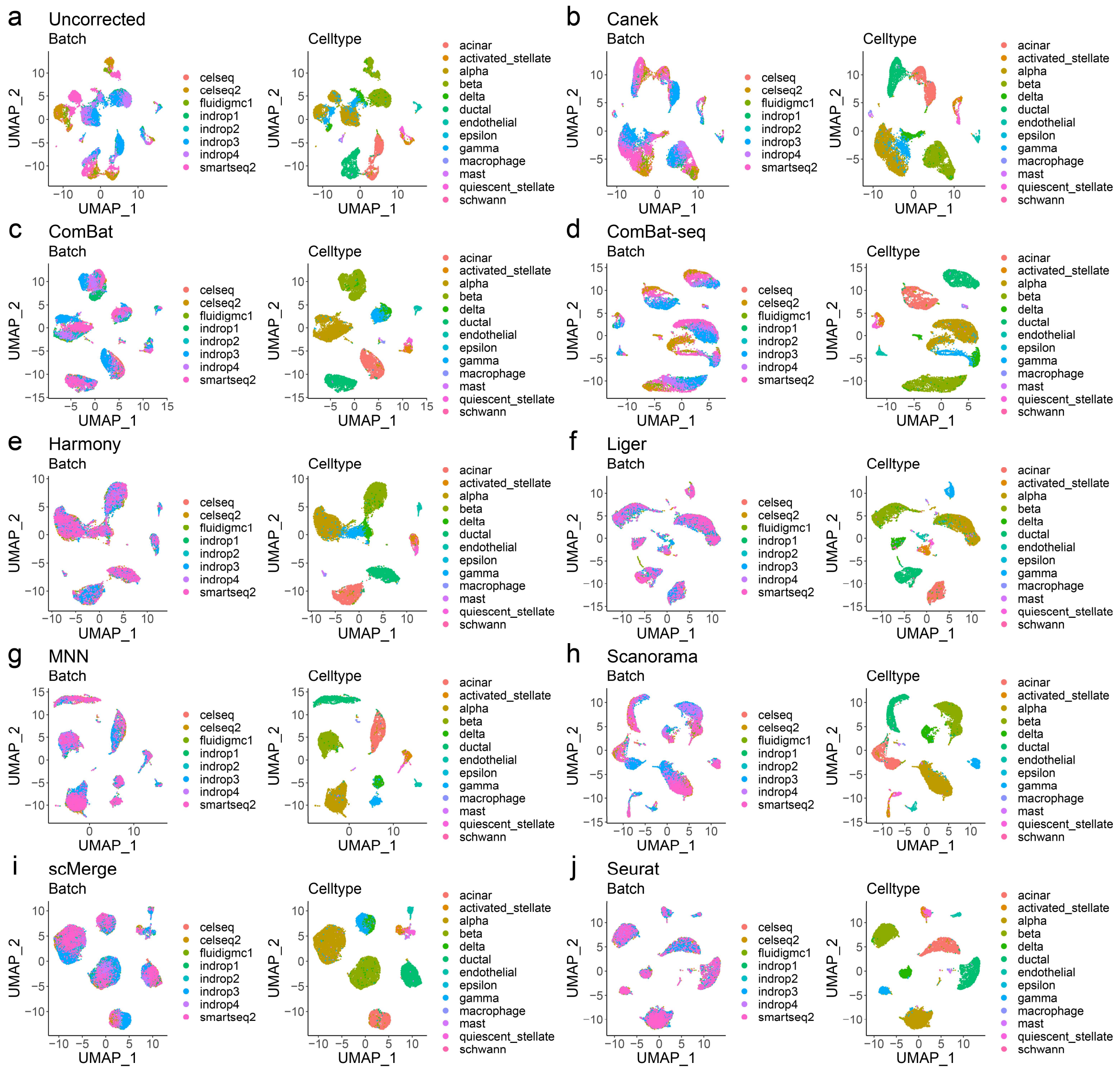


*Supplementary Figure 7. Integration pancreatic islet datasets. Cells are highlighted by cell type.*

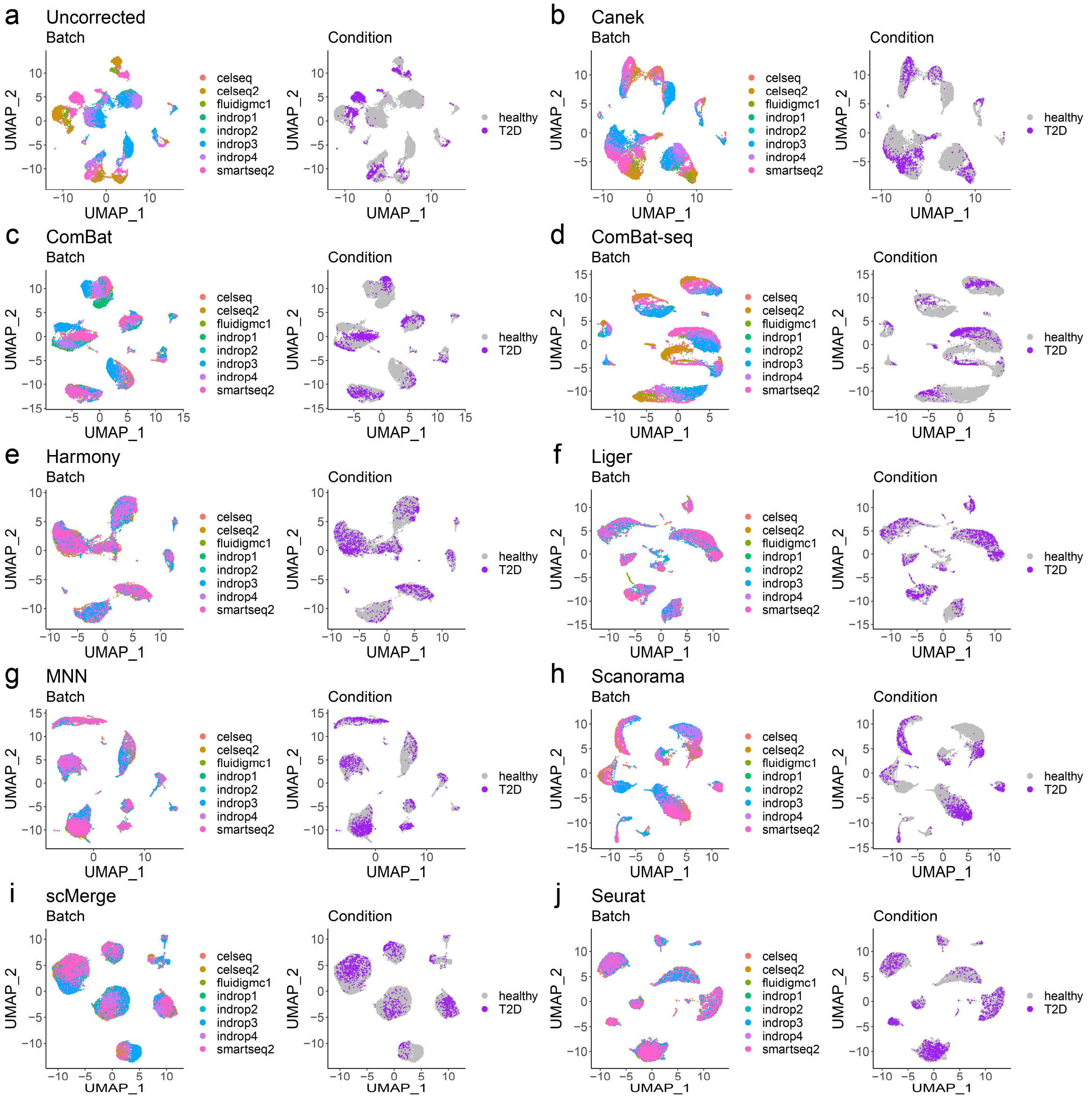


*Supplementary Figure 8. Integration pancreatic islet datasets. Cells are highlighted by disease condition.*

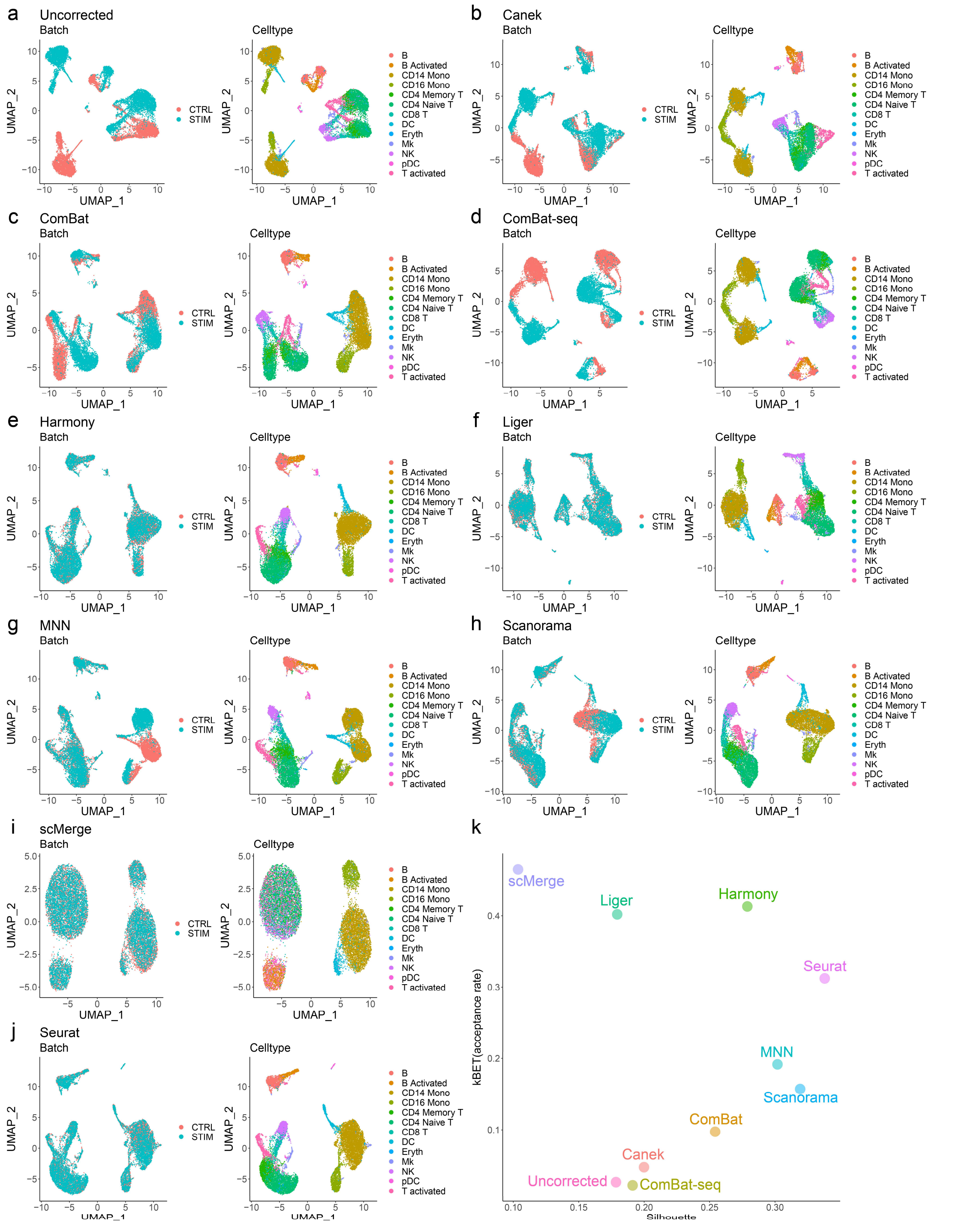


*Supplementary Figure 9. Integration IFN-beta stimulation. a-j) UMAP plots of Uncorrected and after integration. Cells are highlighted by disease condition. k) kBET and Silhouette metrics.*

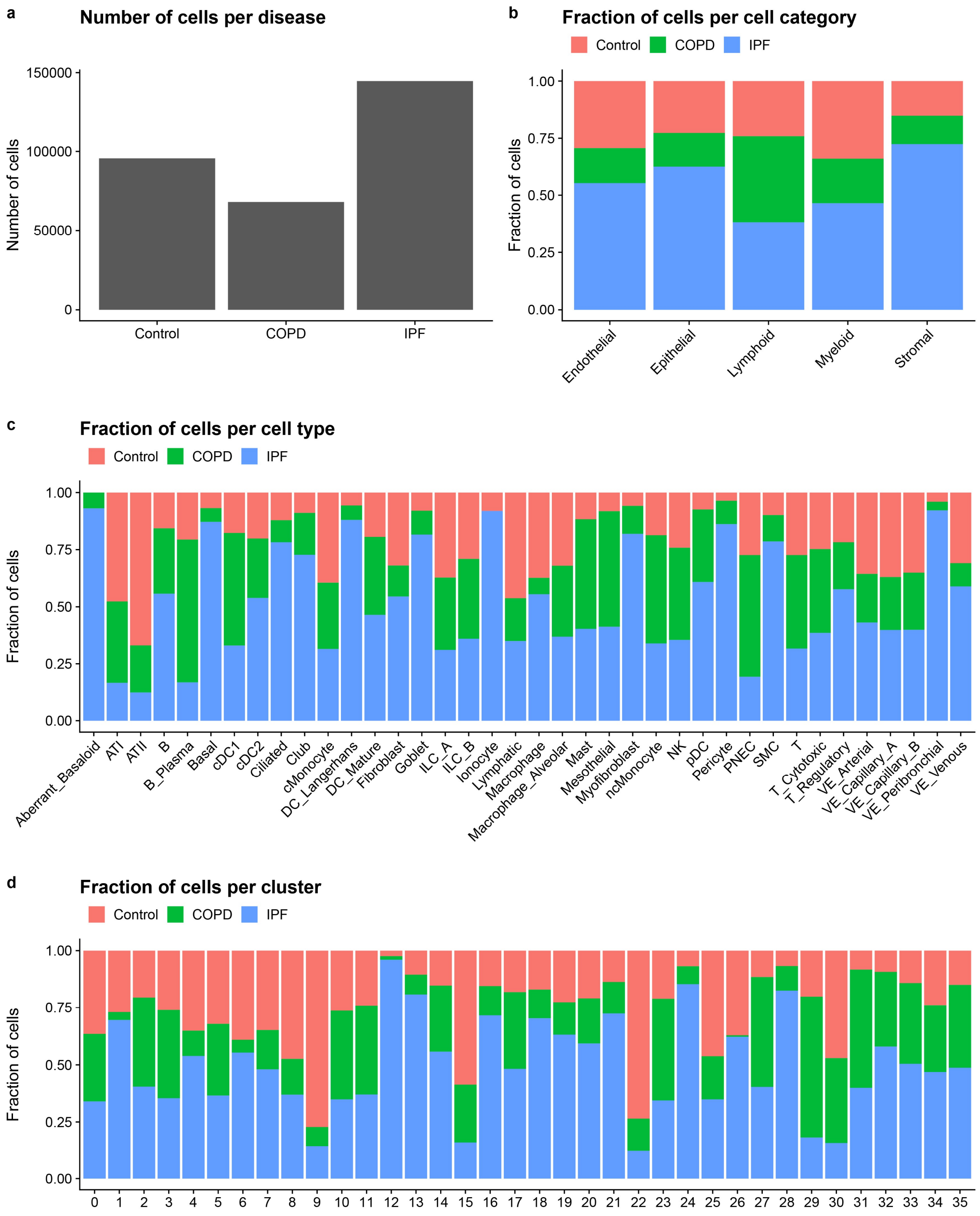


*Supplementary Figure 10. Enrichment of cell populations. a) Number of cells per disease. b) Fraction of disease cells per cell type category. c) Fraction of disease cells per cell populations, captures changes in cell populations in agreement with those reported in the original publication (Adams et al., Science, 2020). For example, there is enrichment of airway epithelial cells (i.e, Basa, Ciliated, and Goblet cells) and depletion of alveolar epithelial cells (i.e., ATI, AT2) in IPF donors. Likewise, the reported Aberrant_Basaloid,cells are enriched in IPF. d) Fraction of disease cells per cluster. This shows that enrichment of IPF cells in specific clusters of interstitial and alveolar macrophages (e.g., clusters 12 and 16).*

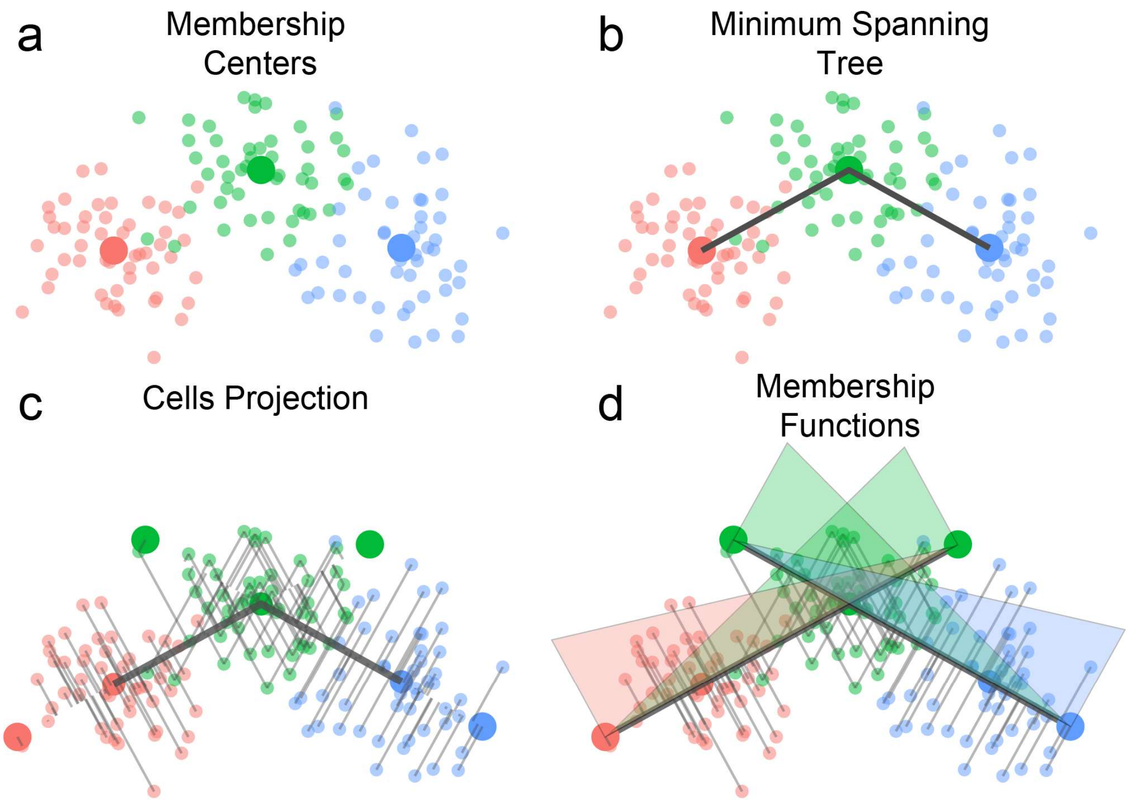


*Supplementary Figure 11. Fuzzy workflow.* ***(a-b)*** *We construct a Minimum Spanning Tree (MST) using the membership center points as nodes.* ***(c)*** *For each edge, we obtain the scalar projection of each of the cells that belong to the memberships and find the maximum and minimum scalars.* ***(d)*** *Using the maximum and minimum scalars we construct a pair of membership functions (MF) for each edge of the MST. The membership functions fuzzy score each of the cells.*

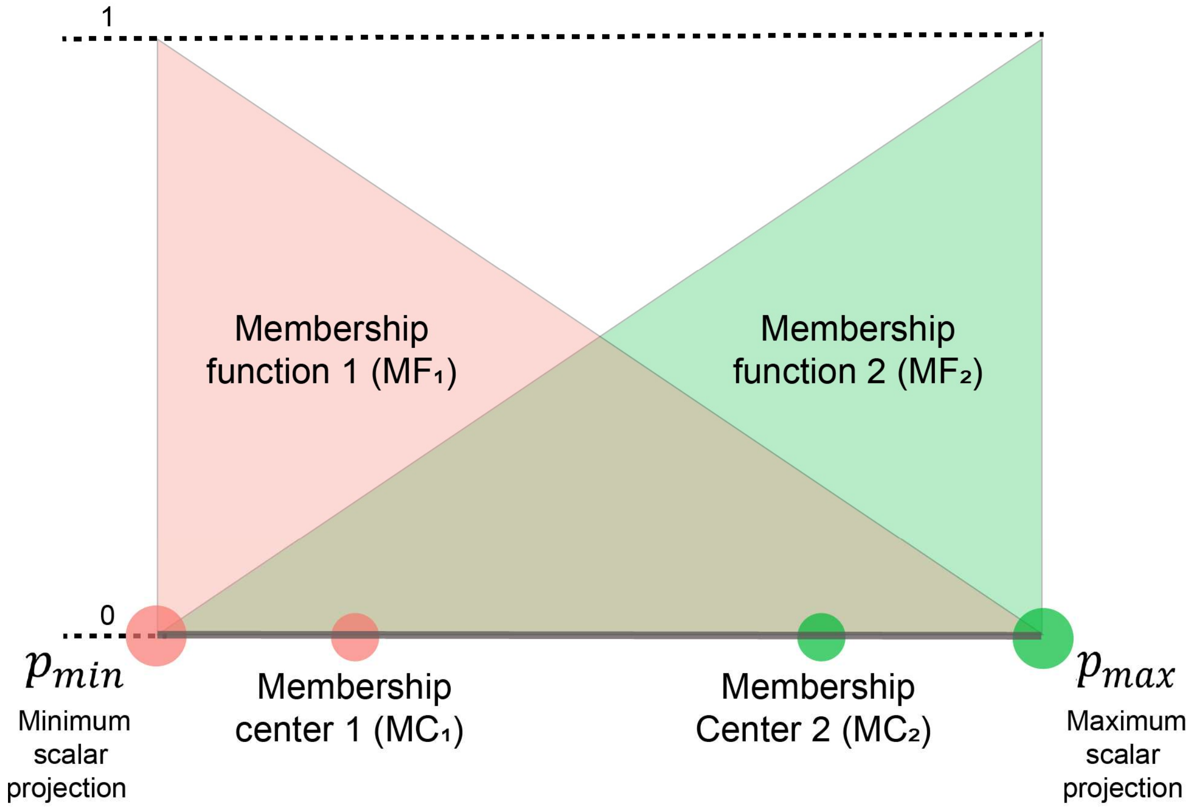


*Supplementary Figure 12. Construction of a pair of membership functions (MFs). The MFs serve as nonlinear mappings from a membership-specific correction to a cell-specific correction. For a given edge, using the vector that joins the membership centers 1 and 2, a scalar projection is found for each of the cells from the two memberships. The MF1 and MF2 are defined such that their minimum and maximum values corresponds with the minimum and maximum scalar projections, and their intersection is located at the midpoint of the line that joins these scalar projections.*
